# Functionally, structurally, and evolutionarily distinct set of genes linked to phenome wide variation in maize

**DOI:** 10.1101/534503

**Authors:** Zhikai Liang, Yumou Qiu, James C. Schnable

## Abstract

In many eukaryotic species the organismal functions of only a small fraction of annotated genes are supported by individual genetic characterization. The organismal functions of a somewhat larger, but still strict minority, of gene models are supported by quantitative genetic analyses (e.g. GWAS). However the organismal functions of the vast majority of gene models are not supported by any direct evidence. Genes characterized by direct investigation exhibit a set of molecular, structural, population genetic, and evolutionary features which distinguish these genes from other gene models. Weaker versions of the same signatures are present among genes identified through conventional quantitative genetics approaches. A new multi-trait multi-SNP association test, the Genome-Phenome Wide Association Study (GPWAS) combines data from large sets of traits and dense resequencing data to identify that set of genes significantly associated with phenotypic variation *per se*. Genes identified using GPWAS and data for 260 phenotypic traits scored across a maize (*Zea mays*) exhibit many of the same molecular, structural, population genetic, and evolutionary signals indicative of genes with functions characterized by direct genetic investigation. The strength of these signals is significantly higher for genes identified using GPWAS than genes identified through conventional GWAS. These results were consistent with a large subset of annotated gene models in maize play little or no role in determining organismal phenotypes. GPWAS and future similar analytical approaches that leverage data from multiple correlated and uncorrelated traits across the same population may provide a method to prioritize those genes most involved in regulation phenotypic variation across diverse species.

## Introduction

In multicellular eukaryotes, only a small proportion of all annotated genes have yet to be linked to loss of function phenotypes^?, 1–6^. Even in extensively studied single celled eukaryotes such as fission yeast (*Schizosaccharomyces pombe*) where the number of annotated genes is less and it is easier to screen for fitness effects under many environmental conditions there are thousands of genes where predicted gene functions have yet to be supported by a loss of function phenotype^7^. The functions of many other genes have been inferred from quantitative genetic analyses. Arguably, the first such quantitative genetic association was the identification of a seed size QTL in dry bean (*Phaseolus vulgaris*) in 1923. This study used a single genetic marker, which was a qualitative trait controlled by a single gene^8^. Soon after, quantitative trait variation could be linked directly to chromosome structural markers^9^. Technology for scoring genetic markers continued to advance, making it possible to genotype markers covering the entire genome across a population. This enabled Genome Wide Association Studies (GWAS) employing the linkage disequilibrium (LD) present in natural populations to identify functionally variable alleles of a gene influencing variation in a target trait^10–12^.

The vast majority of loss of function mutations affect multiple traits. However, the majority of current quantitative genetics approaches seek to identify either genetic markers or genes linked to single phenotypes, with a subset considering data from multiple correlated phenotypes^13–20^. It is now feasible to collect data for thousands of intermediate molecular phenotypes, such as transcript, protein, or metabolite abundance, from entire association populations and incorporate these data into quantitative genetics models as either explanatory^21, 22^ or response variables^23–26^. Advances in high-throughput plant phenotyping have expanded the capacity of these techniques to score dozens or hundreds of whole-organism phenotypes across multiple time points and environments^27, 28^. Incorporating data on large sets of phenotypes scored in the same populations, including both correlated and uncorrelated traits, may aid in the identification of genes that, like the vast majority of classical loss of function mutants, play roles in controlling variation in multiple traits within an organism.

Here we employ a published dataset of 260 distinctly scored traits for 277 resequenced maize inbred lines^29, 30^ to develop and evaluate a novel approach to identify the links between genes and quantitative phenotypic variation *per se* using a multi-trait multi-SNP framework. We demonstrate that the genes identified using this method, which we call Genome-Phenome Wide Association Study (GPWAS), show substantially greater cross-validation in an independent study using data from approximately 20 times as many individuals^31^ than do genes identified using conventional GWAS analysis of the same dataset. For a wide range of features, including expression level and breadth, syntenic conservation, purifying selection in related species, and the prevalence of presence-absence variation (PAV) across diverse maize lines, the genes identified using this multi-trait multi-SNP approach appear more similar to genes identified using forward mutagenesis, and less similar to the overall population of annotated maize gene models.

## Results

Genetic marker data were obtained from resequencing data of 277 inbred lines from the Buckler-Goodman maize association panel^29^. These lines are part of Maize HapMap3, which contains data for a total of 81,687,392 SNPs^30^. After removing the SNPs with high levels of missing data, those that were not polymorphic among the 277 individuals employed here, and several other quality filtering parameters, 12,411,408 SNPs remained. Of these, 1,904,057 SNPs were assigned to 32,084 annotated gene models from the B73 RefGenV4 genome release. Filtering to eliminate redundancy between SNPs assigned to the same gene in high LD with each other reduced this number to 557,968 highly informative SNPs. A phenotypic dataset consisting of 57 specific traits scored for the Buckler-Goodman maize association panel across 1 to 16 distinct environments for a total of 285 unique phenotypic datasets was obtained from Panzea^32^. Removing datasets with extremely high levels of missing data resulted in 260 trait datasets with a median missing data rate of 18%. Of the total 72,020 potential trait datapoints (277 inbred lines × 260 traits), 23.6% or 16,963 trait datapoints were unobserved. Unobserved trait datapoints were imputed using a kinship-based method^33^, and the estimated imputation accuracies for the individual traits are reported in Supplementary Table 1. A conventional GWAS analysis generally employs either empirically determined statistical significance cutoffs^31^, or a Bonferroni correction based on the total number of hypothesis tests^34^ or the number of “effective number” *M*_*eff*_ of independent hypothesis tests conducted^35^. For the above dataset, employing a naive Bonferroni correction would mean each individual analysis would be conducted using a multiple-testing corrected p-value cutoff of 8.96e–08, while a sequential analysis of all 260 traits should employ a multiple-testing corrected p-value of 3.45e–10. As shown in Figure 1a, a given gene might be identified in multiple independent GWAS analyses for individual traits but not be considered significantly associated with any traits when correcting for the total number of traits analyzed. In the example given, Zm00001d002175 shows a statistically significant association with flowering time in multiple environments, yet none of these associations are individually significant enough to meet the threshold for the full multiple testing correction.

**Figure 1.**
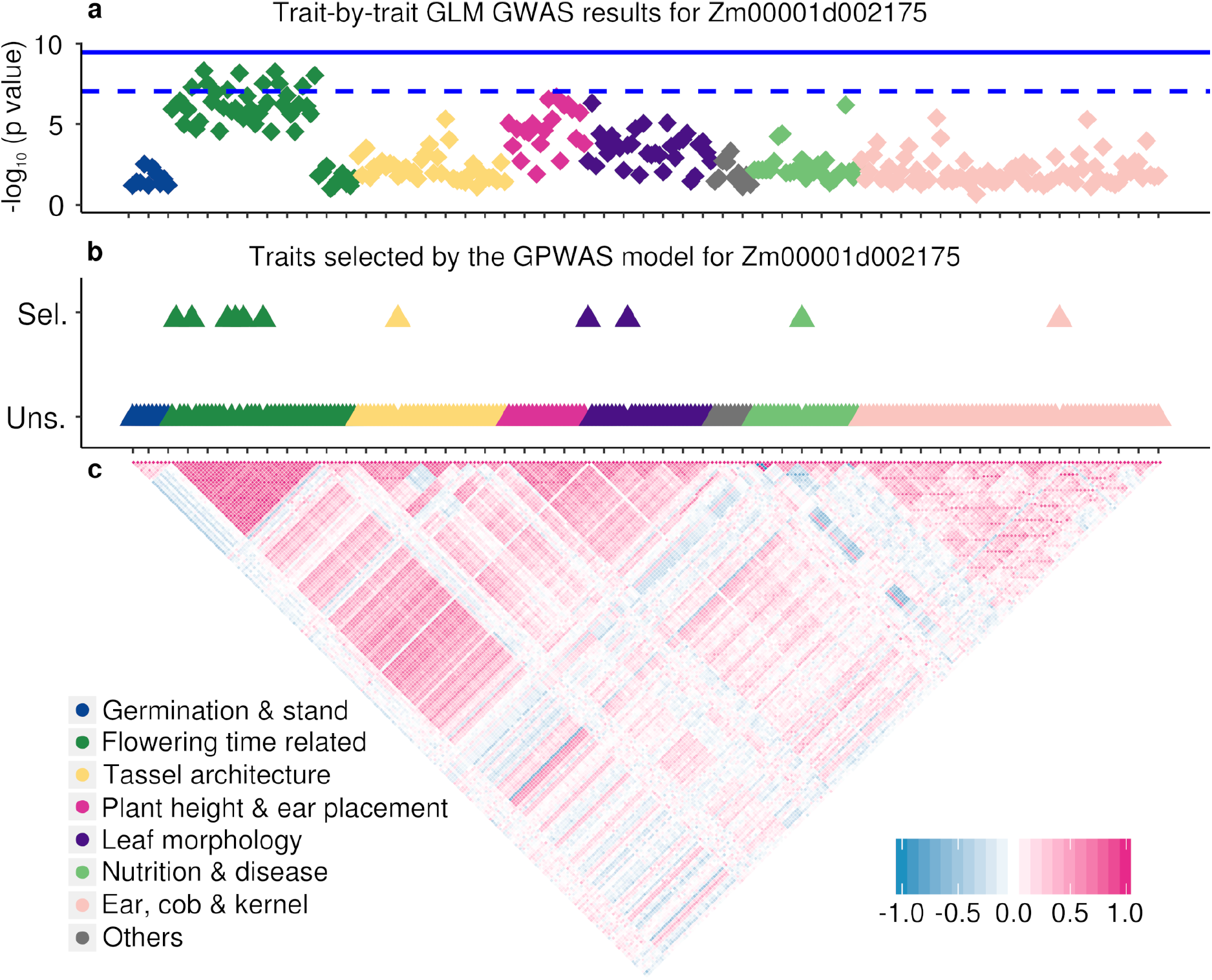
Statistically association between the maize gene Zm00001d002175 and 260 distinct phenotypes. Each diamond or triangle represents one specific phenotypic dataset. Symbol colors indicate the broad categories into which each specific phenotype falls. The specific identities of each phenotype ordered from left to right are given in Supplementary Table 1. (a) The position of each diamond on the y-axis indicates the negative log10 p-value of the most statistically significant SNP assigned to that gene in a GLM GWAS analysis for that single trait. The dashed blue line indicates a p = 0.05 cutoff after Bonferroni correction for multiple testing based on the number of statistical tests in a single GWAS analysis (8.96e–8). The solid line indicates a p = 0.05 cutoff after Bonferroni correction for multiple testing based on the number of statistical tests in GWAS for all 260 traits (3.45e–10). (b) The placement of each triangle on the y-axis indicates whether a given phenotype was included in (Sel.) or excluded from (Uns.) the final GPWAS model constructed for this gene. The complete list of phenotypes incorporated into the GPWAS model for Zm00001d002175 is as follows: days to silk (Summer 2006, Cayuga, NY; Summer 2007, Johnston, NC), days to tassel (Summer 2007, Johnston, NC; Summer 2008, Cayuga, NY), GDD (Growing Degree Days) day to silk (Summer 2006, Cayuga, NY; Summer 2007, Johnston, NC), main spike length (Summer 2006, Johnston, NC), number of leaves (Summer 2008, Cayuga, NY), leaf width (Summer 2006, Champaign, IL), NIR (Near InfraRed)-measured protein (Summer 2006, Johnston, NC) and ear weight (Summer 2006, Champaign, IL). (c) The panel indicates the pairwise Pearson correlation coefficient between each pair of measured phenotypes. Clustering based on phenotypic correlation was used to determine the ordering of phenotypes along the x-axis. Each tick mark on the x-axes of the top and middle panels indicates a distance of five phenotype datasets.

Bonferroni multiple testing correction assumes that each test is independent of all other tests, however, the different trait datasets collected from the Buckler-Goodman association panel exhibited significant correlation (Figure 1c), including three large blocks of traits related to flowering time, plant architectural traits, and tassel structure traits respectively. To address the challenges of partially correlated traits and partially correlated genotype matrices, we developed an approach based upon a stepwise regression model fitting. In this model, the SNPs inside a gene body region are treated as response variables, and both population structure and individual trait datasets are employed to explain the patterns of SNP variance across the population. The significance of the association between each gene and the population of plant phenotypes is determined through a comparison of the phenotype incorporating model with the reduced model incorporating only the population structure variables (see Methods and Supplementary Figure S1).

Multiple testing was corrected using a permutation-based method (see Methods), which controls for the complexities introduced by iterative model selection. Although computationally expensive, permutation has been shown to be robust for controlling false positives in both GWAS and PheWAS studies^36, 37^. Based on the permutation analysis, a p-value cutoff of 1.00e–23 resulted in the classification of 1,776 genes as being significantly associated with phenomic variation in the Buckler-Goodman association panel, resulting in an estimated false discovery rate (FDR) *<* 1.00e-3. Imputation accuracy varied significant among individual traits. To test whether it might be beneficial to exclude traits with high missing data rates and low imputation accuracy, the effect of adding additional simulated zero-heritability phenotypes to the trait matrix was evaluated. Log transformed p-values for individual genes were highly correlated (*R*^2^ = 0.96) before and after the addition of simulated zero heritability traits. All 260 traits were retained for downstream analysis as low information content traits may still provide some value and, based on the test above, appear to be, at worst, benign. For comparison purposes, the same set of traits and genotypes was also tested for associations using three conventional GWAS algorithms: a general linear model (GLM GWAS)^38^, a mixed linear model (MLM GWAS)^38, 39^, and FarmCPU GWAS^40^ (See Methods). Applying an equivalent permutation based FDR threshold to each conventional GWAS algorithm removed the vast majority of positive signals (Supplementary Figure S2). Therefore, for GWAS models, a conventionally multiple testing corrected p-value cutoff was employed (Supplementary Figure S2).

### Validation of Gene-Phenome Associations

A second published dataset of genes identified as being associated with variation in trait values in the maize nested association mapping (NAM) population, which includes approximately 5,000 lines^41^, was employed to assess the relative power and accuracy of three conventional GWAS algorithms as well as the GPWAS algorithm^31, 41^. As the published data for the NAM population used B73 RefGenV2, all comparisons employed only the subset of 29,372 gene models with a clear 1:1 correspondence between gene models included in the B73 RefGenV2 and B73 RefGenV4 annotation versions. Of these, 4,227 of these genes were identified as being associated with at least one trait in the NAM dataset^31^. Genes identified using GPWAS showed significantly higher cross-validation in the NAM dataset than the sets of genes identified using GLM GWAS (p = 2.05e–5; Chi-squared test; two-sided), MLM GWAS (p = 0.010; Chi-squared test; two-sided), or FarmCPU GWAS (p = 0.013; Chi-squared test; two-sided) (Supplementary Figure S3; Supplementary Table 2). Filtering to remove signals from rare SNPs where the minor allele was present in only one or two of the NAM population founder lines reduced the total number of genes identified in that study to 3,621. However, the overall trend observed remained consistent and statistically significant, with the genes identified using the GPWAS algorithm continuing to show statistically significantly higher rates of identification in the reduced NAM dataset (GLM GWAS, p=1.63e-4; MLM GWAS, p=0.002; FarmCPU GWAS, p=0.025; Chi-squared test; two-sided) (Supplementary Table 2). Analyses with two smaller real-world datasets for biochemical traits related to vitamin A (24 traits) and vitamin E (20 traits) metabolism^25, 42^ did not reveal any significant increase in the number of *a priori* gene candidates identified as showing a link to phenotypic variation relative to conventional GWAS approaches (Supplementary Figure S4). Conventional GWAS showed substantial advantages in power at low false discovery thresholds when compared to GPWAS for a single trait in simulation studies, while GPWAS showed significant advantages in power FDR trade offs when data on multiple traits (e.g more than 5 traits) was integrated into the analysis (Supplementary Figure S5). These simulations likely overstate GPWAS’s performance advantage as phenotypes are likely to exhibit different degrees of genetic architecture complexity and differing degrees of shared vs unique causal loci.

Our GPWAS algorithm also produces a list of the specific traits included in the model for a given gene (Supplementary Table 3). For example, in Figure 1b, the overall association between Zm00001d002175 and the trait dataset was statistically significant. The 11 individual traits included in the Zm00001d002175 model included both flowering time measured in multiple locations, as well as additional traits with indirect links to flowering time (e.g. number of leaves, Summer 2008, Cayuga, NY), and others with no obvious links to flowering time. These included the total kernel volume in one year in one location and kernel proteins as estimated using near infrared imaging in another year in a different location.

### GPWAS Accurately Predicts Pleiotropic Consequences of Gene Knockouts

It is important to keep in mind that the associations of individual phenotypes identified within the model are not rigorously controlled for false discovery. We therefore sought to qualitatively evaluate whether traits included in the model for an individual gene make sense in the context of existing detailed biological knowledge about the function of a given gene. One such gene was *anther ear1* (*an1*), a classical maize gene encoding an ent-copalyl diphosphate synthase involved in gibberellic acid biosynthesis, for which knockout alleles have been shown to reduce or abolish tassel branching, reduce plant height, delay growth, and delay flowering^43^. In a separate analysis of the 5,000 individual maize NAM lines, *an1* was identified as being associated with one trait, tassel spike length^44^, however, it was not found to be associated with any individual traits through a conventional GWAS analysis of the Buckler-Goodman 282 dataset. GPWAS identified a statistically significant link between *an1* and a model incorporating multiple phenotypes including flowering time, plant height, and tassel branch number, all consistent with the known mutant phenotypes (Supplementary Figure S6). At least one additional phenotype included in the GPWAS model – germination count (Summer 2006, Johnston, NC) – was not supported by direct reports of characterization of the *an1* knockout allele, but is consistent with the role of *an1* in gibberellic acid metabolism^45, 46^. Overall, the set of phenotypes identified using GPWAS for the *an1* gene appeared to be consistent with previously reports based on either the characterization of the knockout allele or quantitative genetic analyses of natural populations. To disambiguate the effects the multiple SNP and multiple traits portions of this analysis, the same multiple marker analysis for *an1* was conducted separately for each individual trait from the set of 260. This approach identified a larger total number of traits than a joint analysis of all 260 traits together. However, individually significantly associated traits tended to represent a smaller number of phenotype groups (i.e. multiple correlated measurements of the same phenotype in different environments) and failure to capture some of the traits consistent with the known function and mutant phenotypes of *an1* (Supplementary Figure S7).

The GPWAS model also identified *liguleless2* (*lg2*), another classical maize mutant with a well characterized knockout mutant phenotype^47^. The *lg2* encodes a bZIP transcription factor^48^. The loss of *lg2* function disrupts the establishment of the ligule and auricle of the maize leaf and results in plants with extremely erect leaves^47, 49^. Lines carrying *lg2* knockout alleles have been reported to exhibit substantially (10-50%) higher grain yield than otherwise isogenic hybrids^50, 51^, reduced tassel branch numbers^51, 52^, and moderately increased central spike length^52^. Quantitative genetic analyses have identified signals for leaf angle, tassel branch number, and kernel row number associated with the *lg2* locus^36, 44, 53^, although the effect on kernel row number was not significant in at least one study utilizing null alleles of *lg2*^52^. In this study, GPWAS identified a statistically significant link between *lg2* and a model incorporating multiple phenotypes including upper leaf angle, leaf length, central spike length, kernel weight (a yield component trait), and cob diameter. Cob diameter exhibits substantial correlation and overlapping genetic architecture with kernel row number^54^ (Supplementary Figure S8). The GPWAS model for *lg2* also incorporated a number of flowering-time related traits, which do not have consistent support in either the characterization of *lg2* knockout mutants, or previous quantitative genetic analyses of flowering time in maize. Despite this, knockout alleles of *lg2* have been reported to alter the vegetative-to-reproductive phase transition in maize and produce increased numbers of leaves on the main stalk, which would be consistent with its altered flowering time^52^. As in the case of *an1*, the traits identified as being associated with *lg2* using GPWAS appear to be largely consistent with previous characterization of the functional roles of *lg2* in maize.

Individual case studies such as the ones presented above can be misleading. As a control, we set out to identify similar case studies for genes identified by conventional GWAS. A total of seven classical maize mutants from the list in^1^ were identified as linked to one more traits by one of the three GWAS models tested: GLM 5, MLM 1, FarmCPU 1 (Supplementary Table 4). In many cases there was no apparently link between the known gene function and the trait where a significant association was identified. For example, a gene – *glossy8* – involved in the reduction of ketones as part of the biosynthesis of cuticle waxes showed a statistical association with plant tillering, and a transcription factor regulating the production of anthocyanin – *colored alurone1* – showed a statistical association with cob diameter (Supplementary Table 4)^55, 56^. Modestly more promisingly a mutant involved in sex determination – *indeterminate spikelet1/Tasselseed6* – showed a link to flowering time and a second gene involved in cuticular wax wax production – *glossy1* – showed a link to a disease resistance phenotype (Supplementary Table 4)^57, 58^. However, we were not able to identify any stronger links between GWAS trait associations and classical maize mutants in this dataset.

### Greater Functional Specificity of Genes Identified Using GPWAS

Genes identified using GPWAS appear to be a significantly less random sample of total gene models than the set of genes identified using GLM GWAS. A set of 1,406 genes were uniquely identified using GPWAS but not GLM GWAS. An equivalent set of 1,630 genes were identified using GLM GWAS but not GPWAS. In the larger unique-to-GLM GWAS gene set, a single Gene Ontology (GO) term showed a statistically significant bias towards being associated with phenotypic variation (GO:0046034: ATP metabolic process), and two GO terms with nearly identical gene assignments showed a statistically significant bias towards not being associated with phenotypic variation (GO:0000723: Telomere maintenance and GO:003220 Telomere organization). However, the moderately smaller set of genes uniquely identified using GPWAS was enriched or purified for the presence of many more GO terms. A total of 71 GO terms were overrepresented in the unique-to-GPWAS (relative to GLM GWAS) gene set to a statistically significant degree, including numerous terms linked to development, hormone signalling, response to different stimuli, and cell growth (Supplementary Table 5). The 13 GO terms that were underrepresented among genes uniquely identified using the GPWAS algorithm were generally associated with DNA conformation and replication (Supplementary Table 5). A similar comparison was made between genes uniquely identified using GPWAS and FarmCPU GWAS. In this case only 706 genes were uniquely identified using FarmCPU. As it is more likely for an enrichment or purification to be statistically significant in larger populations, only the 706 most significant unique-to-GPWAS (relative to FarmCPU GWAS) genes were evaluated in this comparison to eliminate any potential bias. Among the unique-to-FarmCPU GWAS gene set, only a single GO term was overrepresented to a statistically significant degree (GO:0051707: Response to other organism). However, among the the unique-to-GPWAS (relative to FarmCPU GWAS) gene set of equal size, 39 GO terms showed a statistically significant overrepresentation, while another 4 were statistically underrepresented (Supplementary Table 5).

Several potential factors could explain the large difference in GO enrichment purification we observed between genes identified solely using GWAS and genes identified solely using GPWAS. A number of factors, including the number of GO terms per gene and the proportion of genes with no assigned GO term, differed modestly between the different populations of genes (Supplementary Table 6). The specificity of GO terms varied somewhat between the two populations. The median GO term assigned to a gene identified only using GLM GWAS was assigned to 514 total distinct gene models in B73 RefGenV4. For genes identified only using GPWAS, this decreased to 430 gene models. This difference in the number of genes that a given GO term is assigned does not appear to explain the differences observed in the enrichment or purification (Supplementary Figure S9). Rather, the large differences observed here are consistent with GWAS identifying a more random subset of annotated genes as being associated with phenotypic variation than did GPWAS.

### Molecular, Structural, and Evolutionary Features of Genes Identified Using GPWAS

Genes identified using the GPWAS algorithm differed from the overall population of annotated maize gene models in a number of characteristics, as well as from the populations of genes identified using conventional GWAS. In many cases, the properties of genes identified using GPWAS appeared more similar to the population of genes with validated loss-of-function phenotypes^1^. Slightly less than half of all annotated maize genes were expressed to a level above 1 fragment per kilobase of transcript per million mapped reads (FPKM) on the average of the 92 tissues/time points assayed^59^. This figure was greater than 2/3 for the genes identified using the three conventional GWAS algorithms, and approximately 3/4 for genes identified using the GPWAS algorithm and maize genes with validated loss-of-function phenotypes (Figure 2a; Supplementary Table 2). Genes identified using GLM GWAS, MLM GWAS, FarmCPU GWAS, GPWAS, and the classical mutants all exhibited greater breadths of expression across tissues, larger numbers of genes with observed evidence of translation, and greater gene lengths than the population of annotated genes as a whole (Supplementary Table 2; Supplementary Table 7). The number of associated SNPs was positively correlated with the log-transformed inverse p-value assigned to genes using both GWAS (r = 0.566) and GPWAS (r = 0.625) (Supplementary Figure S10; Supplementary Table 7). However, this association declined dramatically in the permuted data for GPWAS (median permuted r = 0.155), but remained high for GWAS (median permuted r = 0.626) (Supplementary Table 8). This suggests that the high number of SNPs per gene for GPWAS (median: 43 SNPs, mean: 47.3 SNPs) relative to the overall gene set (median: 12 SNPs, mean: 17.4 SNPs) is a biological property of the genes controlling phenotypic variation in this population, rather than reflecting a bias in the GPWAS algorithm.

**Figure 2.**
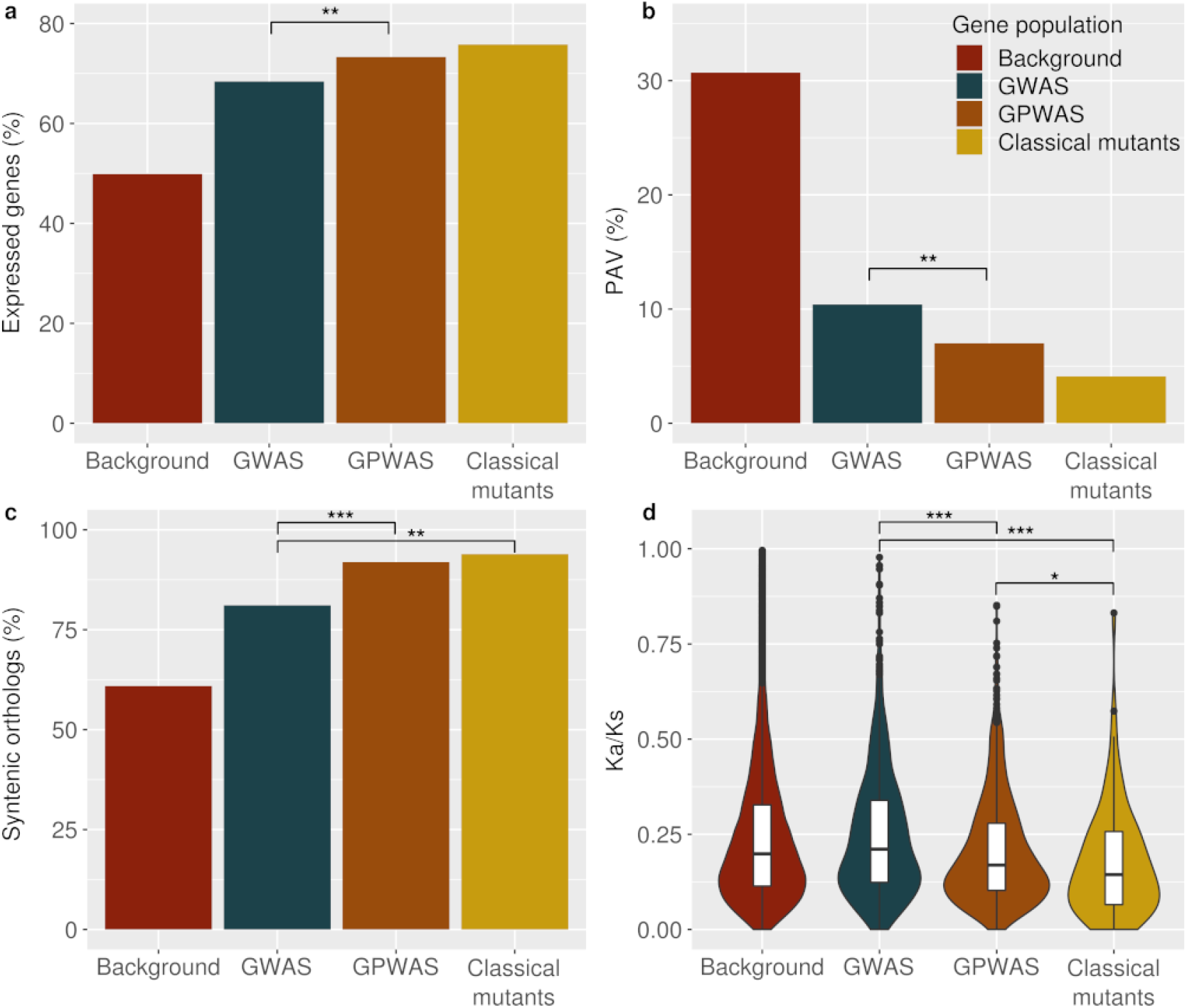
Comparisons among four gene populations: Background, GWAS (Genes linked to phenotype by GWAS (GLM) but not by GPWAS), GPWAS (Genes linked to phenotype by GPWAS but not by GWAS (GLM)) and classical mutants with known loss of function phenotypes^1^. (a) Proportion of genes within each of the four populations which express with average FPKM *>* 1 of 92 assayed tissues/time points. (b) Proportion of genes within each of the four populations which exhibit presence absence variation (PAV) in maize. (c) Proportion of genes within each of the four populations which are conserved at syntenic orthologous locations in sorghum (*Sorghum bicolor*). (d) Distribution of non-synonymous substitution rate/Synonymous substitution rate rations (Ka/Ks) for the subset of genes from each of the four populations with syntenic orthologs in both sorghum and foxtail millet (*Setaria italica*). A small number of individual genes with Ka/Ks ratios *>*1 are not show to aid readability (one gene in the GLM GWAS population, two in the GPWAS population). *:p value *<* 0.05; **: p value *<* 0.01; ***: p value *<* 1e-3 (Chi-Squared test employed for panels a-c; Mann–Whitney U employed for panel d.). GWAS, GPWAS, and classical mutant populations exhibited statistically significant differences from the background gene population for all four features shown. Comparable data for the other two GWAS algorithms evaluated as part of this study (MLM and FarmCPU), are provided in Supplementary Table 9.

On a population and comparative genomics level, genes identified using the GPWAS algorithm also differed from the overall population of annotated maize gene models, and looked more like genes with validated loss-of-function phenotypes. Genes identified using both the conventional GWAS and GPWAS algorithms were significantly less likely to exhibit PAV in the maize populations (Figure 2b) than the overall population of maize gene models. The reduction in PAV frequency for genes identified using GPWAS (7.0%) was statistically significantly greater than for genes identified only using GWAS (10.4%) (p=0.0015; Chi-squared test; two-sided), and not statistically significantly different from low level of presence absence variation observed for maize genes with validated loss of function phenotypes genes (4.1%) (p=0.36; Chi-squared test; two-sided) (Supplementary Table 9). Genes identified using either conventional GWAS and GPWAS algorithms were significantly more likely to be conserved at syntenic orthologous locations in sorghum than the overall set of maize gene models (Figure 2c). Genes uniquely identified using GPWAS were more likely to be conserved at syntenic locations in the genome of sorghum *Sorghum bicolor* (91.8%) than those uniquely identified using GWAS (74-85%; see Supplementary Table 9). This difference was statistically significant in comparisons to all three GWAS algorithms tested and was comparable to the likelihood of syntenic conservation for maize genes with known loss of function mutant phenotypes (93.9%) (Supplementary Table 9).

The genes identified as being associated with phenotypic variation using GPWAS also appeared to be under stronger purifying selection than either the overall population of maize gene models or those identified using any of the three conventional GWAS algorithms (Figure 2d; Supplementary Table 9). This analysis was constrained to the subset of gene models with conserved orthologs in sorghum (*Sorghum bicolor*), and foxtail millet (*Setaria italica*). Among these genes, those uniquely identified using GPWAS showed a reduced ratio of nonsynonymous substitution rate to synonymous substitution rate (Ka/Ks) (median: 0.168–0.169; mean 0.208–0.210), relative to the overall population of syntenically conserved maize gene models (median: 0.200; mean: 0.246), while those uniquely identified using GWAS showed elevated rates (median: 0.202–0.233; mean: 0.251–0.261) relative to the same overall population (Supplementary Table 9). Among the maize genes with characterized loss-of-function phenotypes, this ratio declined even further (median: 0.144; mean: 0.177). In short, the typical annotated gene appears to experience notably less purifying selection than those associated with organismal-level phenotypic variation based on either characterized loss-of-function mutant phenotypes or those identified using the GPWAS, but not a GWAS, algorithm.

## Discussion

Complex datasets can contain scores for dozens or hundreds of traits across the same populations. The prevalence of these datasets and the challenges and opportunities they present is expected to grow in the coming years. Here, we developed an approach for identifying genotype-phenotype associations that can scale to the analysis of datasets containing hundreds, or potentially even thousands, of traits. The statistical tests upon which the GPWAS approach is built become unstable once the number of traits exceeds the number of individuals scored, therefore, scaling to high numbers of traits would require the use of larger association populations than many of the most general used plant populations today^10, 29, 60, 61^. Multicollinearity in either the predictor or response variables can make the statistical estimation and inference procedures we employed unstable^62^. One common approach for reducing the total number of traits in a multi-year and/or multi-field site trial is to calculate the best linear unbiased predictors (BLUPs), which provide a single value for a given trait in a given line across multiple environments^63^. However, this approach strips out information on trait plasticity across environments, controlled by distinct sets of genes from those controlling multiple environment mean values^64^ and is thus likely to bias the downstream analysis away from a large class of genes involved in determining organismal phenotypes across changing environments. In cases where the number of measured traits exceeds the number of environments, it would be advisable to employ alternative approaches to reduce the dimensionality of the trait dataset, whether that be an *ad hoc* approach such as selecting a subset of representative traits from highly correlated blocks, or dimensional reduction analyses such as a principal component analysis or multidimensional scaling. The automatic application of variable selection and/or dimensional reduction in such scenarios could be incorporated into future GPWAS implementations.

Conventional approaches to multivariate GWAS analysis require that the set of traits being tested simultaneously be correlated^19, 20^. These algorithms cannot scale efficiently to testing associations with hundreds of traits simultaneously on a genome-wide scale, as adding more traits in multivariate GWAS leads to exponential increases in computational cost^13^. GPWAS can iteratively search through large sets of correlated and uncorrelated traits. While this provides advantages in terms of scalability, it does come at some cost. Firstly, as presently implemented, GPWAS cannot estimate marker/gene effects as is possible with many conventional GWAS algorithms. Secondly, as a result of the stepwise selection procedure, the set of traits identified as associated with a gene in a GPWAS analysis is unlikely to be exhaustive, particularly when multiple traits are closely correlated. For example, tassel spike length and tassel length are generally correlated, including in this dataset. GPWAS identified variation in tassel length but not spike length as associated with *an1*. However, if tassel length is removed from the dataset, spike length becomes one of the phenotypes selected by the GPWAS model for *an1*

Another challenge for the present implementation of GPWAS is that it requires regions of interest to be defined across the genome. In this study, annotated gene models were used to define these regions, however, approximately 40% of the phenotypic variation in maize has been estimated to be explained by noncoding regulatory regions^65^. These regions can be separated from the genes whose expression they control by many kilobases^66, 67^, while LD in maize generally decays within one to several kilobases^68, 69^. Both sequence conservation and chromatin mark data could be used to define additional regions of interest likely to represent regulatory sequences^65, 70–74^. Similar approaches could also be employed to identify currently unannotated regions of the genome with a high potential of containing cryptic genes, including functional long noncoding RNAs^75^.

## Conclusion

We found that genes with statistically significant links to phenotypic variation exhibit substantial differences from the overall population of annotated genes in the maize genome for a number of characteristics. They are more likely to be transcribed to significant levels, more likely to be conserved at syntenic orthologous positions in the genomes of related species, dramatically less likely to exhibit PAV across diverse maize inbred lines, and appear to be subjected to notably stronger purifying selection than the overall population of annotated genes. In all these cases, the genes identified using GPWAS are less like the overall population of annotated gene models, and more like the small subset of genes in the maize genome whose functions have been characterized using loss of function alleles^1^. Based on the results and controls presented above, it appears that these differences between genes significantly associated with trait variation and the overall population of annotated gene models in maize reflects a biological reality, rather than an algorithm bias. The distinct features shared by both genes identified using classical forward genetics and now using GPWAS suggest that it is unlikely that all annotated genes in the maize genome contribute to organismal phenotypes. Over the past three decades, without substantial discussion or debate, many in the scientific community have moved from a definition of genes that was based on organismal function, to one which is based on molecular features^76–78^. However, many analyses still implicitly assume that genes annotated in the genome based on homology and/or expression evidence must play a role in determining organismal phenotypes. The absence of evidence for a role in determining a phenotype is interpreted as a failure to find either the correct trait to measure or the correct environment in which to measure it. Improved approaches to distinguish which annotated gene models are more likely to contribute to controlling organismal phenotypes will be critical to future efforts to guide gene-by-gene functional characterization efforts.

## Methods

### Genotype and Phenotype Sources, Filtering, and Imputation

Raw genotype calls from the resequencing of the maize 282 association panel^30^ were retrieved from Panzea in AGPv4 coordinates. Missing genotypes were imputed using Beagle (version: 2018-06-10)^79^. Only biallelic SNPs with fewer than 20% missing data points were subjected to imputation. After imputation, SNPs with a minor allele frequency (MAF) of less than 0.05 or which were scored as heterozygous in more than 10% of samples were discarded. A phenotype file (traitMatrix maize282NAM v15-130212.txt) containing total of 285 traits, corresponding to 57 unique types of phenotypes scored in 1 to 16 environments was downloaded from Panzea. A set of 277 accessions with identical names in the HapMap3 data release and the Panzea trait data were employed for all downstream analyses.

Maize gene regions were extracted from AGPv4.39, which was downloaded from Ensembl. SNPs were clustered based on *R*^2^ *>* 0.8 and only one randomly selected SNP per cluster was retained. If, after collapsing the highly correlated clusters, the number of SNPs exceeded 138 (50% of the number of inbred lines scored), a random subsample of 138 SNPs was employed for the downstream analyses. Identical final SNP sets were employed for the GPWAS and GWAS analyses.

Of the 285 initial trait datasets, 25 were removed because the data file contained a recorded trait value for only one individual, leaving a total of 260 trait datasets. Using a Bayesian multiple-phenotype mixed model^33^, missing phenotypes were imputed based on a kinship matrix calculated from 1.24 million SNPs generated using GEMMA (version: 0.94.1)^17^. For those traits with a sufficient numbers of real observations to enable evaluation, the accuracy of the phenotypic imputation was assessed independently by masking 1% of available records for each trait and comparing the imputed and masked values. This process was repeated 10x for each trait.

### Principal Components Used in GPWAS and GWAS

A subset of 1.24 million SNPs distributed across both intragenic and intergenic regions on all 10 chromosomes was used to perform PCA using the R prcomp function to provide controls for population structure. As the maize 282 panel exhibits relatively low population structure^29^, only the first three PCs were included in both GWAS and GPWAS analyses, however, comparable analyses could be run with different numbers of PCs included. In GPWAS, for analysis of the given gene on each chromosome, markers solely from the other 9 chromosomes were used to reduce the endogenous correlations between genes and principal components^80, 81^. In GWAS, principal components were calculated using all of 1.24 millon SNPs on 10 chromosomes.

### GPWAS Analysis

All the operations for the GPWAS analyses are detailed in the R source code used to conduct the analysis – and associated documentation – which has been made available online (https://github.com/shanwai1234/GPWAS). Briefly, we employed a model selection approach to adaptively select the most significant phenotypes associated with each gene. A F-test (one-sided) was used to compare a model to explain variation in SNPs based solely on population and a model which incorporated both population structure and trait data. The significance in the difference of the goodness of fit between these two models was used to determine the significance of the association of individual genes with phenotypic variation in the dataset, the Reduced Model (RM) based solely on population structure and the Phenotype incorporating Model (PM) which used stepwise selection to include both population structure and trait data within its model.

#### Reduced Model

Let the subscripts *k* and *i* represent the *k*th individual and the *i*th gene. Let *g*_*k,i*_ be the corresponding SNP values in that gene. Here, *g*_*k,i*_ is an *m*-dimensional vector of SNPs where *m* is the number of distinct genetic markers associated with the gene after removal of SNPs in high linkage disequilibrium with each other. We considered the multiple responses regression model

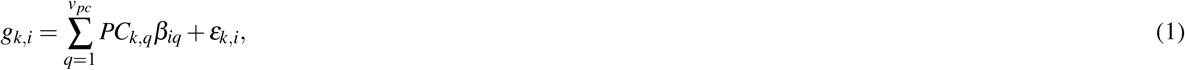

where *q* denotes the *q*th principal component, and *v*_*pc*_ is the number of the included principal components in (1). The regression coefficients *β*_*iq*_ and the errors *ε*_*k,i*_ are all *m* dimensional vectors.

#### Stepwise Selection

The final model begins with the initial model (1) and uses a stepwise selection procedure^82^ to add additional traits as explanatory variables. In each iteration, we consider the model

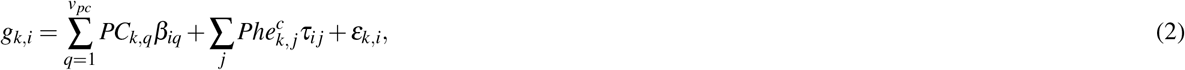

where 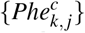 are the currently selected phenotypes, and *τ*_*ij*_ is the corresponding coefficient for the phenotype 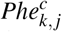 of the *i*th gene. The goodness of fit for each trait in the model (2) was assessed for all SNP markers in the *i*th gene jointly.

In detail, least square estimation was applied to fit the regression (2) and to obtain the residuals 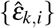. Let 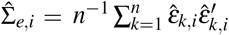 be the residual sample covariance. The association between the *k*th trait and all of the evaluated SNPs were jointly evaluated by comparing the determinants of residual sample covariances 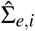 with and without that trait included in the model. The p-value of the association was obtained from a F-test built on a likelihood ratio statistic widely used in multi-response regression (see Section 7.7 in Johnson et al. (2002)^83^). This F-test incorporated the dependence among the SNPs, and thus provided a more powerful test than the individual test of association for each single SNP by combining multiple signals across different SNPs.

If at least one trait passed a set significance threshold – here p *<* 0.01 was selected – the single most significant trait among all traits significantly associated at p *<* 0.01 was added to the model (2); see Supplementary Figure S1. The model itself was then rerun using all traits selected to that point. If any of the traits already incorporated into the model failed to meet the original cut-off value of p *<* 0.01 after the incorporation of the newest trait, the single least significantly associated trait was removed from the model Supplementary Figure S1.

The process above constituted one iteration of the stepwise selection procedure. In the analyses presented in this manuscript, 35 sequential iterations of the stepwise selection procedure were performed per gene. With this dataset of genotypes and traits, every gene tested converged to a single stable model within less than 35 iterations (Supplementary Figure S11), however this assumption would need to be revisited when employing GPWAS on other datasets which might include either more individuals, more traits, or fewer traits with high pairwise correlation.

The cut-off value 0.01 is widely used for stepwise regression^82^. Looser thresholds (.05 and 0.1) required significantly more iterations to converge, dramatically increasing computational cost. The final threshold for statistical significance of the overall model was determined using a permutation based method, as described below, limiting the potential for this cut-off value to influence final outcomes.

#### Phenotype incorporating Model

The final phenotype incorporating model (PM) can be represented as:

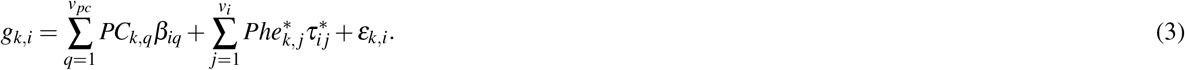

In the final model (3), there were *v*_*i*_ selected phenotypes for the *i*th gene, where *v*_*i*_ ≤ 260. The selected phenotypes 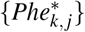 were a subset of the collection of all the phenotypes {*Phe*_*k*,1_, *Phe*_*k*,2_, …, *Phe*_*k*,260_}, and 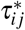 was the corresponding coefficients for the selected phenotype 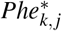 of the *i*th gene. Note that *g*_*k,i*_, *β*_*iq*_, and 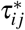 can be vectors corresponding to the multiple SNPs within the *i*th gene.

#### Model Comparison

The final step was to evaluate how much the inclusion of trait data improved model fit (PM) relative to a purely population structure based model (RM). The statistical significance of the increase in goodness of fit of the two models was compared using a F-test^83^ via comparing the residuals covariances of those models. The F-test takes into account all of the SNPs included from the target interval, as well as the degree of correlation between these SNPs. The filtering of highly linked SNPs described above satisfies the criteria of the F-tests that multiple response variables should not exhibit strong correlations with each other.

As adding more explanatory variables to a model will always tend to improve the goodness of fit, permutation based analyses were used to determine a threshold for a statistically significant increase in goodness of fit between the reduced model (RM) and the phenotype incorporating model (PM). Twenty permutations of the trait/genotype associations were conducted and GPWAS was run independently on each of these twenty permutations. Distributions of F-test PM/RM model comparisons from the permuted and unpermuted data were used to estimate false discovery rates at different cut off thresholds (Figure S2).

## GWAS Analysis

GLM GWAS and MLM GWAS analyses were conducted using the algorithm first defined by Price and coworkers^38^. The FarmCPU GWAS with the algorithm was defined by Liu and colleagues^40^. All of algorithms were run using the R-based software rMVP (MVP version 1.0.1) (A Memory-efficient, Visualization-enhanced, and Parallel-accelerated Tool For Genome-Wide Association Study) (https://github.com/XiaoleiLiuBio/rMVP). FarmCPU analysis method was run using maxLoop = 10 and method.bin = “FaST-LMM”^84^. The first three principal components were considered to be additional covariates for the population structure control in all of analyses. The same kinship matrix used in the phenotype imputation was also used for controlling the genotype relationship in the MLM GWAS model, while the method for analyzing variance components (vc.method) was set to GEMMA^85^. To enable a comparison with the GPWAS results, each gene was assigned the p-value of the single most significant SNP among all the SNPs assigned to that gene across the 260 analyzed phenotypes in the GWAS model.

### Classical Mutant Analysis

Maize classical loss of function mutant identities were taken from a previous study^1^. To obtain an exhaustive list of reported mutant phenotypes, papers were mined from MaizeGDB loci pages, where both papers and conference abstracts that reference studies on individual maize genes, both cloned and uncloned, are captured by manual data curation^86^.

### Nested Association Mapping Comparison

Published associations identified for 41 phenotypes scored across ~5,000 maize recombinant inbred lines were retrieved from Panzea (http://cbsusrv04.tc.cornell.edu/users/panzea/download.aspx?filegroupid=14)^31^. Following the thresholding proposed in that paper, a SNP and CNV (copy number variant) hits with a resample model inclusion probability ≥0.05, which were either within the longest annotated transcript for each gene (AGPv2.16) or within 15kb upstream or downstream of the annotated transcription start or stop sites were assigned to that gene respectively. Gene models were converted from the B73 RefGenV2 to B73 RefGenV4 using a conversion list published on MaizeGDB (https://www.maizegdb.org/search/gene/downloadgenexrefs.php?relative=v4).

### Gene Expression Analysis

Raw reads from the a published maize expression atlas generated for the inbred line B73 were downloaded from the NCBI Sequence Read Archive PRJNA171684^59^. Reads were trimmed using Trimmomatic-0.38 with default setting parameters^87^. Trimmed reads were aligned to the maize B73 RefGenV4 reference genome using GSNAP version 2018-03-25^88^. Alignment results were converted to a sorted BAM file format using Samtools 1.6^89^, and the FPKM values were calculated for each gene in the AGPv4.39 maize gene models in each sample using Cufflinks v2.2^90^. Only annotated genes located on 10 maize pseudomolecules were used for downstream analyses and the visualization of the FPKM distribution.

### Ka/Ks Calculations

For each gene listed in a public syntenic gene list^91^, the coding sequence for the single longest transcript per locus was downloaded from Ensembl Plants. Their sequences were each aligned to the single longest transcript of genes annotated as syntenic orthologs in *Sorghum bicolor* v3.1^92^ and *Setaria italica* v2.2^93^, retrieved from Phytozome v12.0 using a codon-based alignment as described previously^94^. The calculation of the ratio of the number of nonsynonymous substitutions per non-synonymous site (Ka) to the number of synonymous substitutions per synonymous site (Ks) was automatically calculated using scripts which are provided on github (https://github.com/shanwai1234/Grass-KaKs). Genes with a synonymous substitution rate less than 0.05 on the branch leading to maize after the maize/sorghum split were excluded from the analyses, as these atypically low Ks values tended to produce extreme Ka/Ks ratios. Genes with multiple tandem duplicates were also excluded from the Ka/Ks calculations. The calculated Ka/Ks ratios of maize genes are provided in Supplementary Data 1.

### Presence/Absence Variation (PAV) Analysis

PAV data were downloaded from a published data file^95^. Following the thresholding proposed in that paper, a gene was considered to exhibit presence absence variance if at least one inbred line had a coverage of less than 0.2.

### Gene Ontology Enrichment Analysis

All GO analyses used the maize-GAMER GO annotations for B73 RefGenV4 gene models^96^. Statistical tests for GO term enrichment and purification were performed using the goatools software package (v0.8.12)^97^, with support for a two-sided Fisher’s exact test provided by the fisher exact function in SciPy. To determine the median information content of the GO term, each was assigned a score based on the total number of gene models to which this GO term was assigned to in the maize-GAMER dataset. This analysis considered only gene models to which a GO term was specifically applied to in the dataset, but not gene models where the assignment of the GO term may have been implied by the assignment of a child GO term. Genes in B73 RefGenV4 Zm00001d.2 that employed in maize-GAMER GO annotations (~40,000 genes) were used as the background population.

### Evaluation of GPWAS and GWAS Power and FDR Using Simulated Data

SNP calls for the entire set of 1,210 individuals included in Maize HapMap3 were retrieved from Panzea^30^, filtered, and assigned to genes as described above resulting in 1,648,398 SNPs assigned to annotated gene body regions in B73 RefGenV4. Two thousand genes, associated with 30,547 SNP markers were randomly sampled for downstream simulation. Independent phenotypes with known causal QTNs were simulated using the additive model in GCTA (v1.91.6)^98^. Effect sizes for each QTN for each simulated phenotype in each permutation were drawn from a normal distribution centered on zero.

The resulting simulated trait data and genuine genotype calls were analyzed using GLM GWAS, FarmCPU GWAS, and GPWAS as described above, with the exception that the population structure PCs were calculated using a sample (1% or 191,856 SNPs) of the total SNPs remaining after filtering, rather than only using the subset of SNPs assigned to the 2,000 randomly selected genes included in this analysis.

For each analysis, the set of 2,000 genes was ranked from most to least statistically significant based on the significance of the most significantly associated SNP (for GLM and FarmCPU GWAS) or the significance of the overall model fit relative to a population structure only model (for GPWAS). The power evaluation for GPWAS was defined as the number of true positive genes relative to the total number of causal genes, and FDR was defined as the number of false positive genes relative to the total number of positive genes. Power and FDR were calculated in a steps of five genes starting with the five most significant genes and continuing to the 500 most significant genes (i.e. {5,10,…,495,500}).

In each simulation, 100 genes (5%) were randomly selected as causal genes. For each causal gene in each simulation, a causal SNP was selected to simulate the phenotypic effects. A total of 100 phenotypic traits with heritability as 0.5 were simulated. Total 100 simulated phenotypes were split into 1, 5, 10, 20, 50 and 100 subgroups for running GPWAS.

## Acknowledgements

This work is supported by National Science Foundation Awards MCB-1838307 and OIA-1826781 to JCS. In addition, we received support from the Quantitative Life Sciences Initiative at the University of Nebraska-Lincoln, which in turn received support from the University of Nebraska Program of Excellence. This work was completed utilizing the Holland Computing Center of the University of Nebraska, which receives support from the Nebraska Research Initiative. The authors thank Andy Dahl for the advice and instruction in the use of phenotype imputation software, Zheng Xu and Wenlong Ren for their consultation on the design of the association study, and the Panzea project (http://www.panzea.org) for gathering the phenotypic and genotypic data employed in this study.

## Additional information

Supplementary Table 1: The 260 phenotypes employed in this study with corresponding missing data rates, imputation accuracies and classified phenotype classes.

Supplementary Table 2: Expression characteristics, protein abundance and NAM gene validation among gene populations.

Supplementary Table 3: Significant genes detected using GPWAS and the phenotypes selected for each gene model.

Supplementary Table 4: Classical maize mutants identified as associated with quantitative trait variation by one or more GWAS algorithms, but not GPWAS.

Supplementary Table 5: GO terms enriched and purified in gene populations uniquely identified in GPWAS.

Supplementary Table 6: Statistics of GO terms assigned to each gene population.

Supplementary Table 7: Gene length and SNP density in each gene population.

Supplementary Table 8: Correlation between significance levels and SNP numbers per gene for the genes generated from permuted and real data in GPWAS and GLM GWAS.

Supplementary Table 9: Conservation features for unique gene sets between each of GWAS models (GLM GWAS, MLM GWAS and FarmCPU GWAS) and GPWAS.

Supplementary Data 1: Categories of annotated maize genes (AGPv4.39).

**Figure S1.**
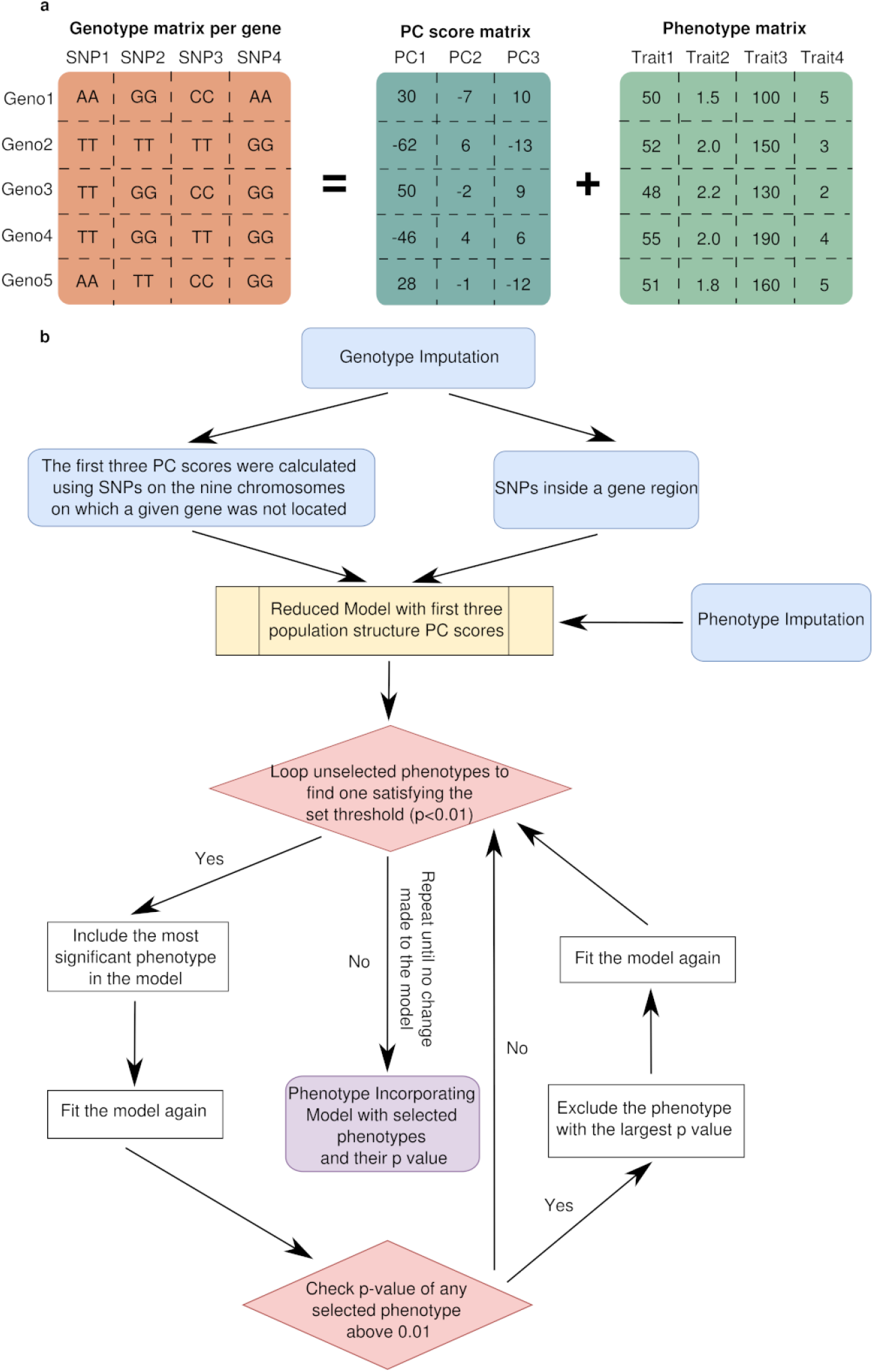
GPWAS algorithm implementation. (a) Example of trait and genotype matrices employed for GPWAS. (b) Flow chart showing initial data processing and the forward selection process within the GPWAS algorithm.

**Figure S2.**
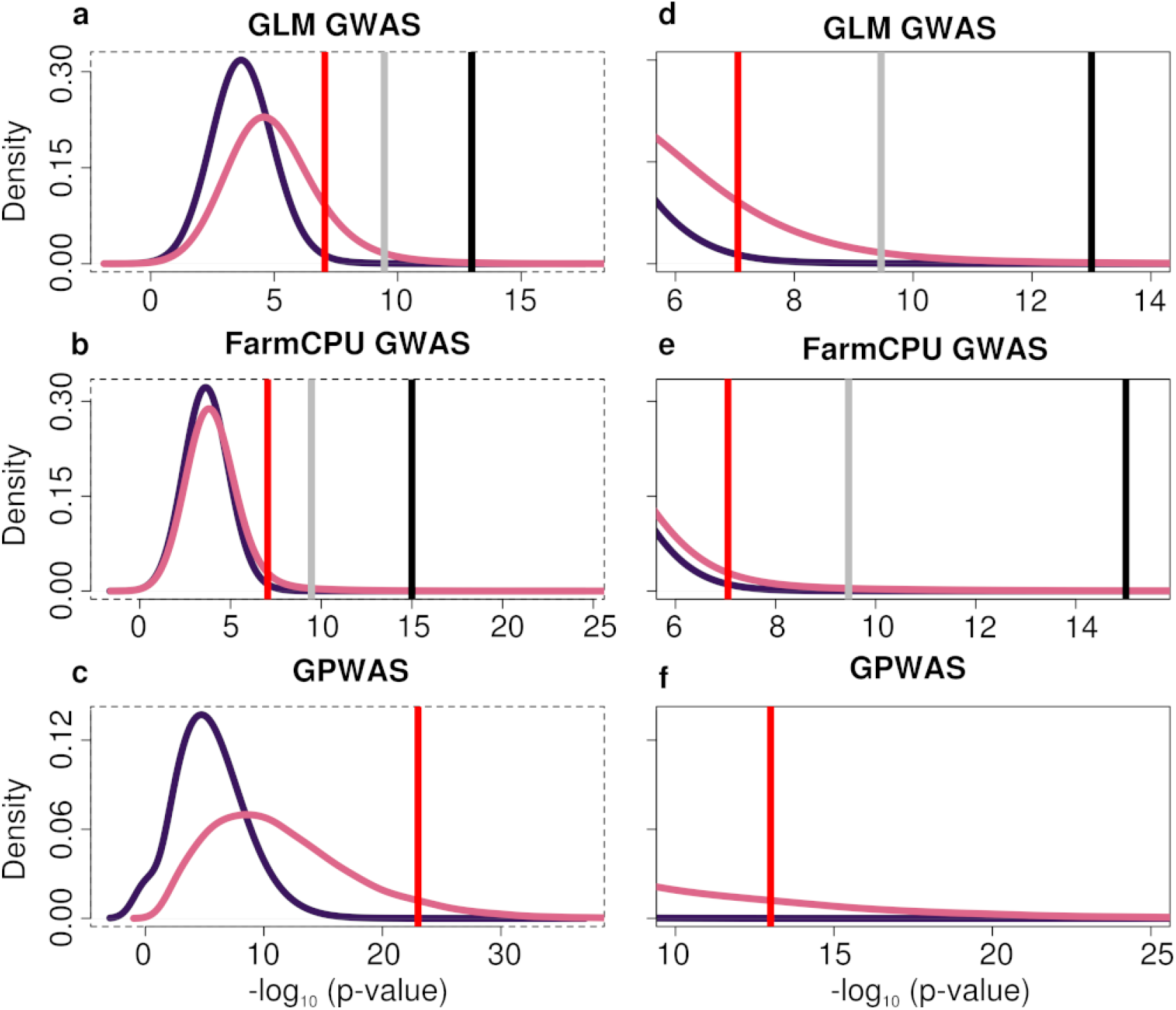
Permutation testing based estimation of false discovery rates for GLM GWAS, FarmCPU, and GPWAS. For each panel, the dark curve shows the distribution of per gene p-values obtained from 20 permutations of genotype and trait data (see Methods), while the light curve indicates the distribution of per gene p-values obtained from the analysis of the non-permuted dataset. Red lines indicate the p-value analyses employed in these analysis, corresponding top p-value = 8.96e-8 for GLM and FarmCPU and an estimated FDR *<* 0.001 for GPWAS. Genes assigned p-values on the right side of each red line were employed for all downstream analyses in the main text. Panels a-c show the entirety of the distributions, while panels d-f display a magnified view of the regions of the curve where the p-value threshold is employed. When these data were used to estimate the p-value cut off corresponding to an estimated FDR *<* 0.001 for GLM GWAS, this was found to correspond to an uncorrected p-value of approximately 1e-14, resulting in 31 genes would remain statistically significantly associated with traits. For FarmCPU GWAS, the minimum FDR achieved was FDR *<* 0.029 at a p-value threshold of 1e-15, resulting in 38 genes remaining statistically significantly associated with traits.

**Figure S3.**
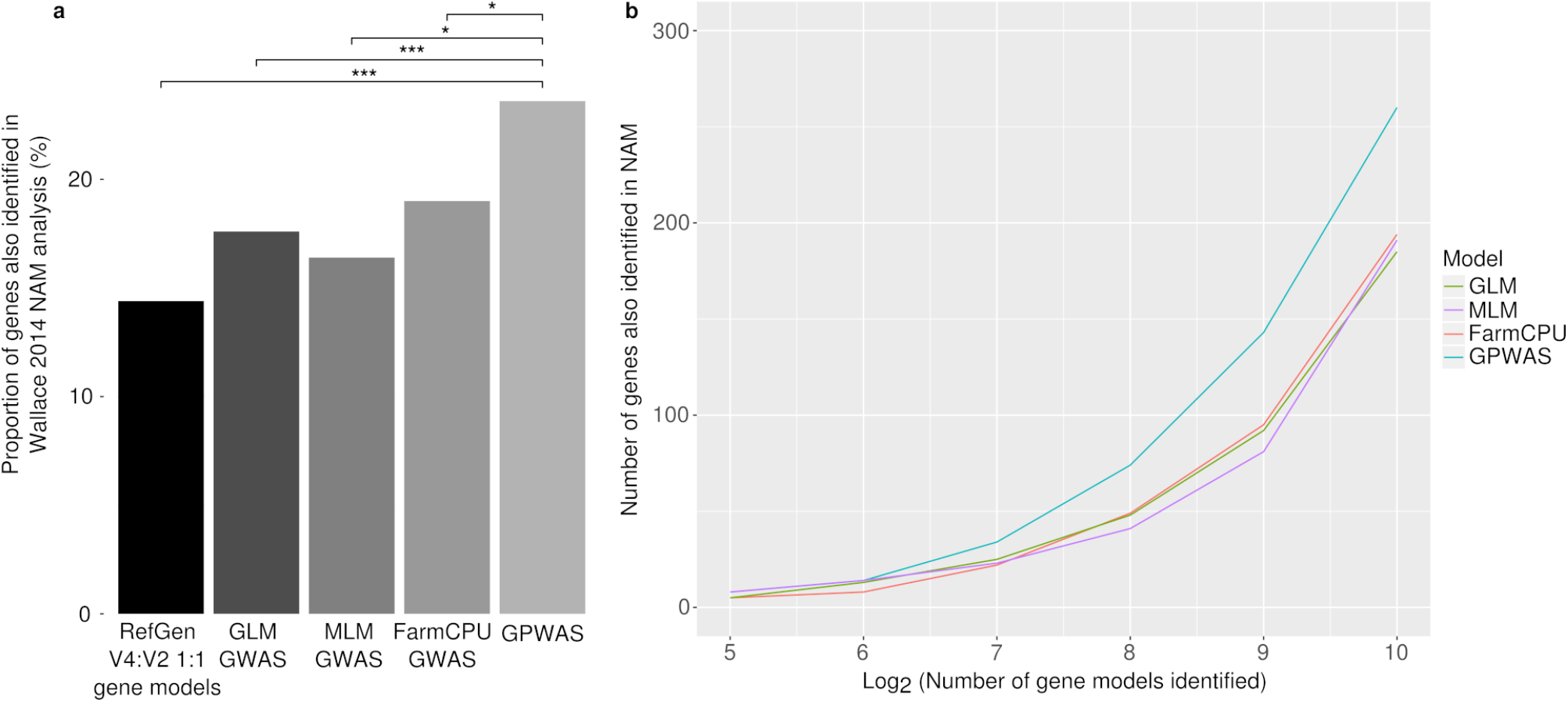
Cross validation of gene candidates identified using either GLM GWAS, MLM GWAS, FarmCPU GWAS, or GPWAS of the maize 282 association panel using an independent set of gene candidates identified in the 5,000 line maize nested association population by Wallace et al^31^. (a) The background rate of genes identified by Wallace et al among all maize gene models with 1:1 relationships between B73 RefGen v2 (used by Wallace et al) and B73 RefGen v4 (used in this study), as well as higher proportions of genes identified by each of the four quantitative genetics methods which also overlap with Wallace et al. *: p value ≤0.05; ***: p value ≤1e-3 (Chi-Squared Test). (b) Relationship between the total number of positive genes selected by each of the four quantitative genetics methods and the total number of positive genes which were also identified by Wallace et al.

**Figure S4.**
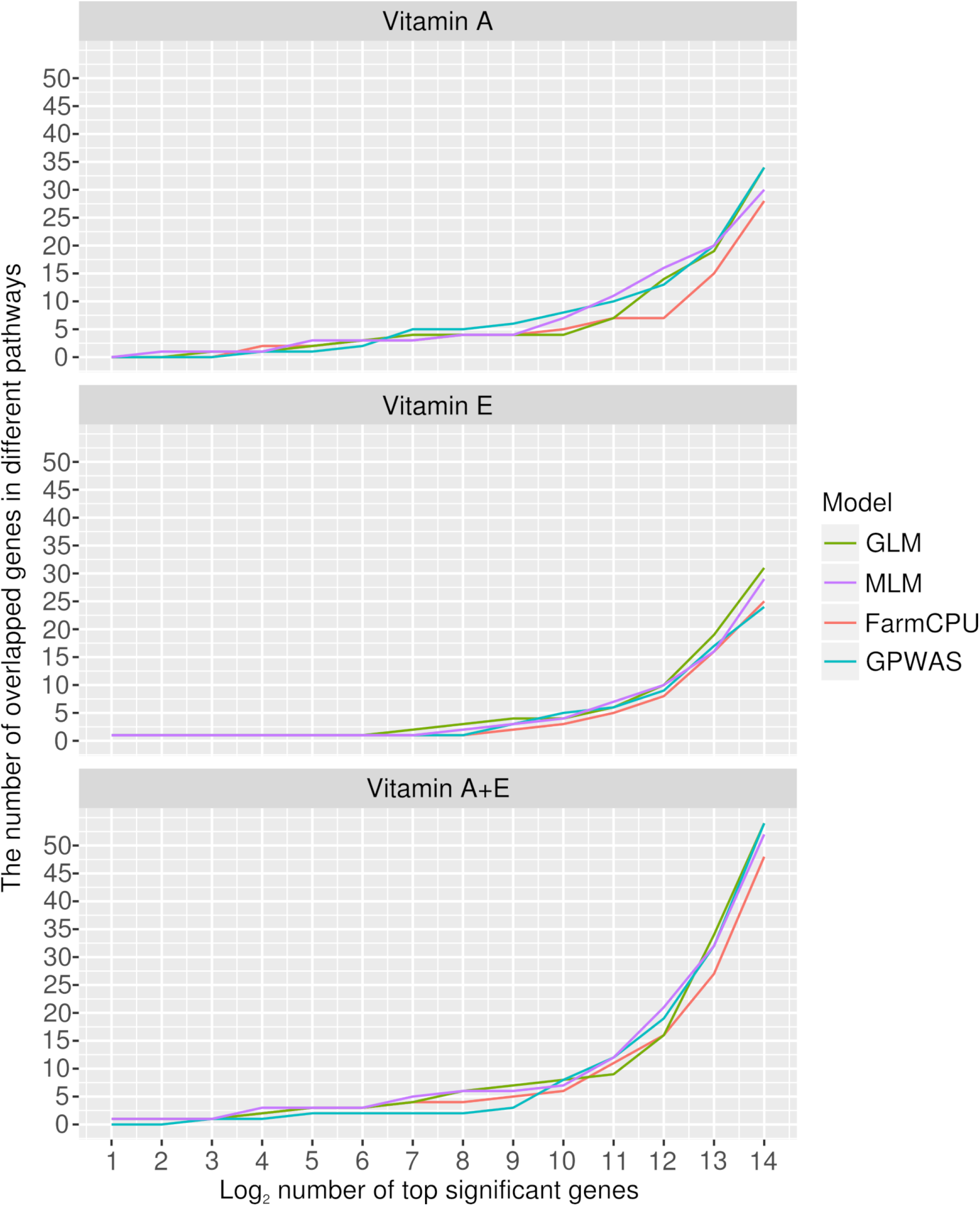
Comparison of the performance of GPWAS and conventional GWAS methods in the identification of *a prior* candidate genes involved in vitamin A and E biosynthesis. Phenotypic data and published *a priori* candidate gene lists for vitamin A and vitamin E were taken from previous studies^25, 42^. The methodology used here was otherwise identical to that employed for Supplementary Figure S3.

**Figure S5.**
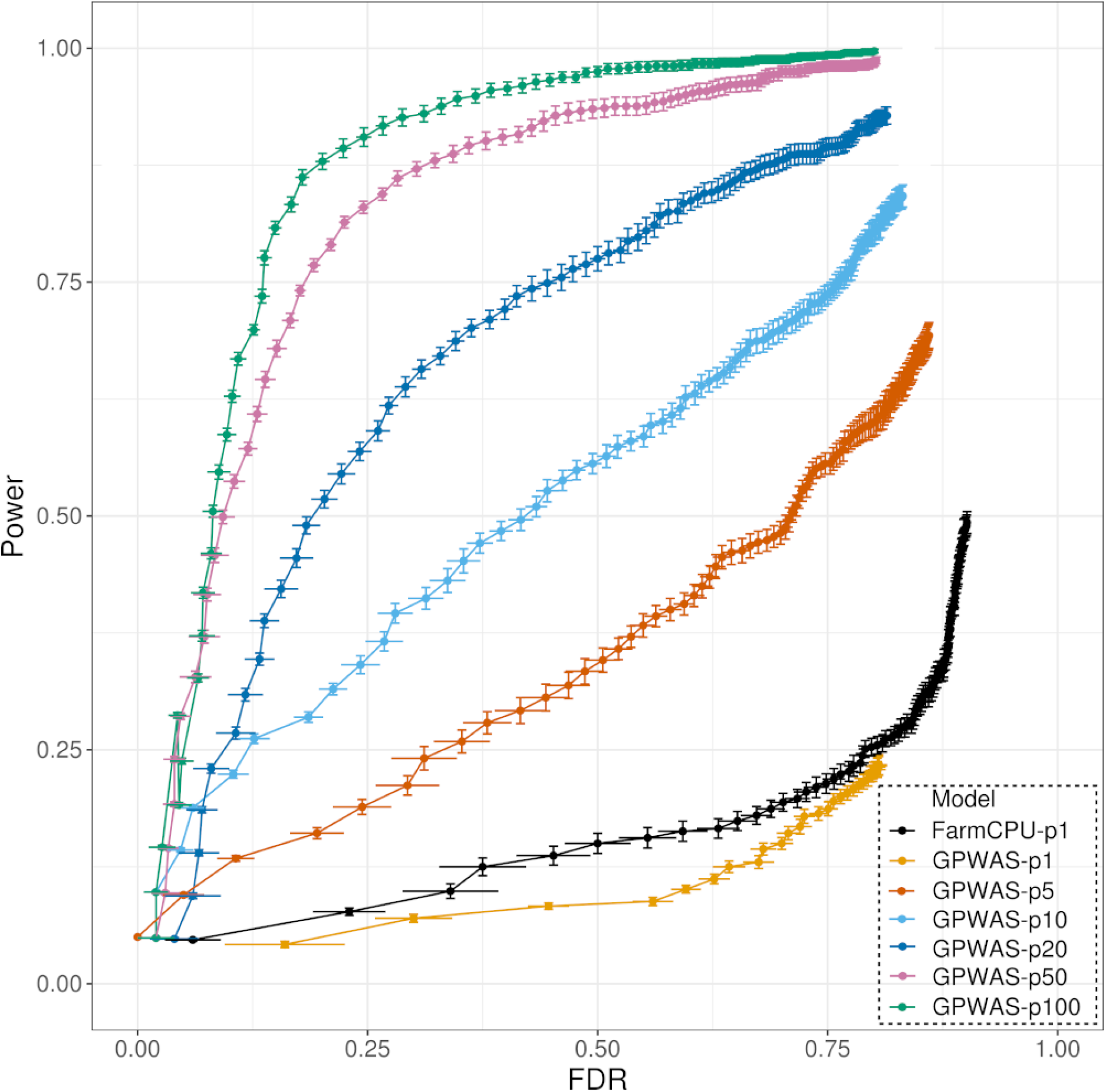
Comparison of power and FDR trade-offs between GPWAS and FarmCPU GWAS when working with simulated data. The comparison shown is based on 10 different simulations, each including data for between 1 and 100 phenotypes controlled by 100 randomly selected quantitative trait nucleotides (QTNs) each with heritability of (h^2^=0.5). FDR and power were calculated in step sizes of 5 positive genes starting with the top 5 most statistically significant genes for each method, then 10, 15,.. 500. Points indicate mean values for FDR and power at each step across the 10 independent simulations and error bars indicate standard errors for both FDR and power. FarmCPU provided more favorable trade offs between power and FDR than the other two GWAS models at all steps and those these two other methods are omitted for readability. GPWAS-p1 indicates results using a single simulated phenotype, GPWAS-p2 indicates results using 2 simulated phenotypes for running GPWAS. The same naming standard is applied to GPWAS-p5, GPWAS-p10, GPWAS-p20, GPWAS-p50 and GPWAS-p100. FarmCPU outperformed GPWAS when both methods used data from a single phenotype. GPWAS was only able to outperform FarmCPU when it was able to incorporate data from multiple phenotypes controlled by pleiotropic genes.

**Figure S6.**
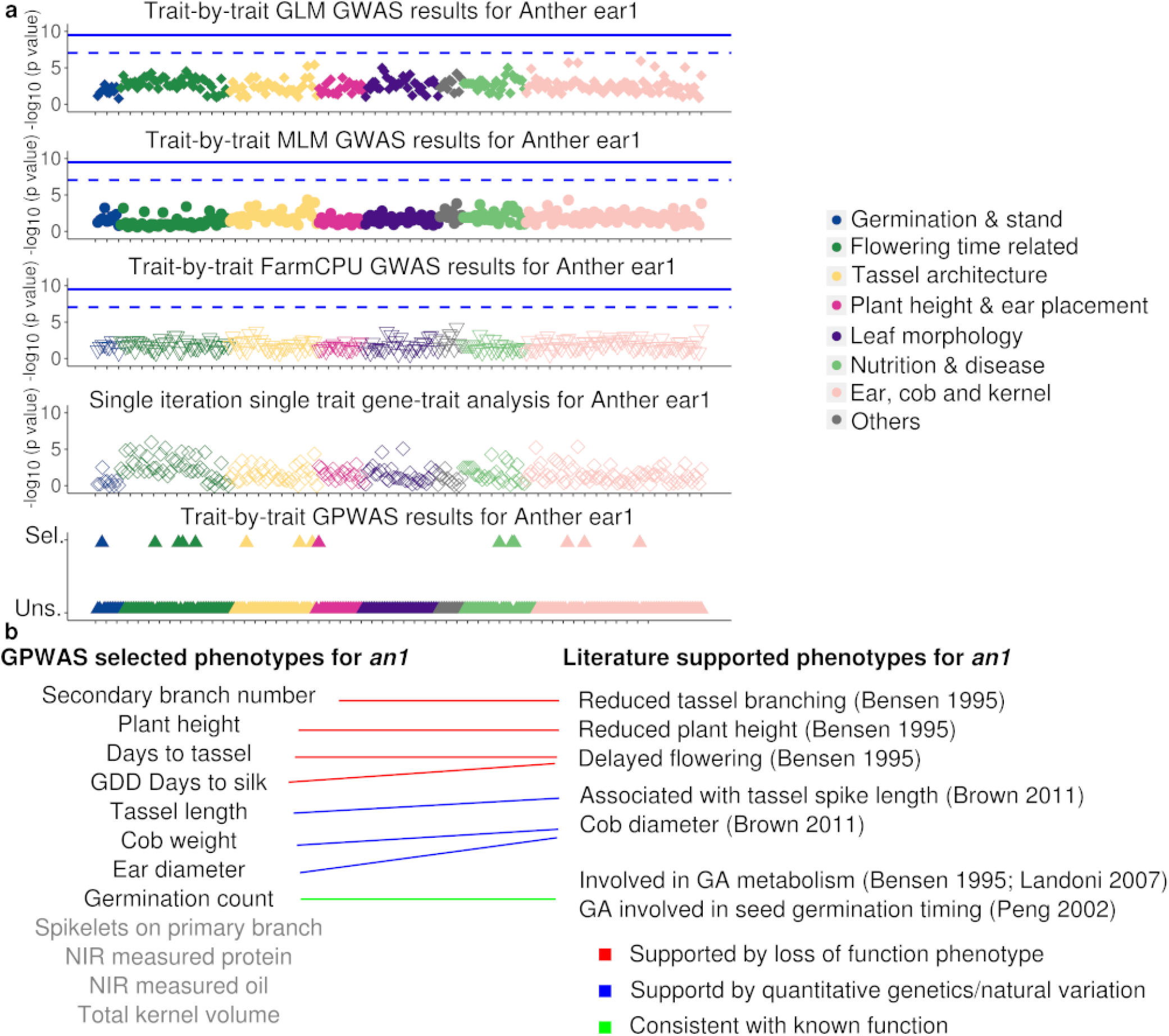
Evaluation of GLM GWAS, MLM GWAS, FarmCPU GWAS, single iteration single trait gene-trait association and GPWAS using the known maize gene *Anther ear1* (*an1*) (Zm00001d032961). (a) The dashed lines indicates a p-value corresponding to 0.05 after a Bonferroni correction for independent tests on 557,968 (SNPs). Solid lines indicate the stricter multiple testing corrected threshold, which considers both the number of SNPs and the number of phenotypes tested. In the GPWAS panel, Sel. and Uns. indicate traits that were selected and unselected respectively, in the model GPWAS fit for this particular gene. Phenotypes are ordered along the x-axis in the same order used for Figure 1, with each tick mark indicating a distance of five phenotypes. Phenotypes incorporated in the GPWAS model for *an1* were as follows: germination count (Summer 2006, Johnston, NC), days to tassel (Summer 2007, Cayuga, NY), GDD days to silk (Summer 2007, Johnston, NC; Summer 2007, Champaign, IL; Winter 2006, Miami-Dade, FL), tassel length (Summer 2007, Cayuga, NY), spikelets primary branch (Summer 2006, Champaign, IL), secondary branch number (Summer 2006, Boone, MO), plant height (Summer 2006, Cayuga, NY), NIR-measured protein (Summer 2006, Johnston, NC), NIR-measured oil (Summer 2006, Johnston, NC; Winter 2006, Miami-Dade, FL), cob weight (Summer 2007, Johnston, NC), ear diameter (Summer 2007, Johnston, NC) and total kernel volume (Summer 2006, Cayuga, NY). (b) The potential correspondence between phenotypes selected using the GPWAS model for *an1* using the GPWAS model and phenotypes either reported for loss of function *an1* mutants or previous quantitative genetic analyses^43–46^.

**Figure S7.**
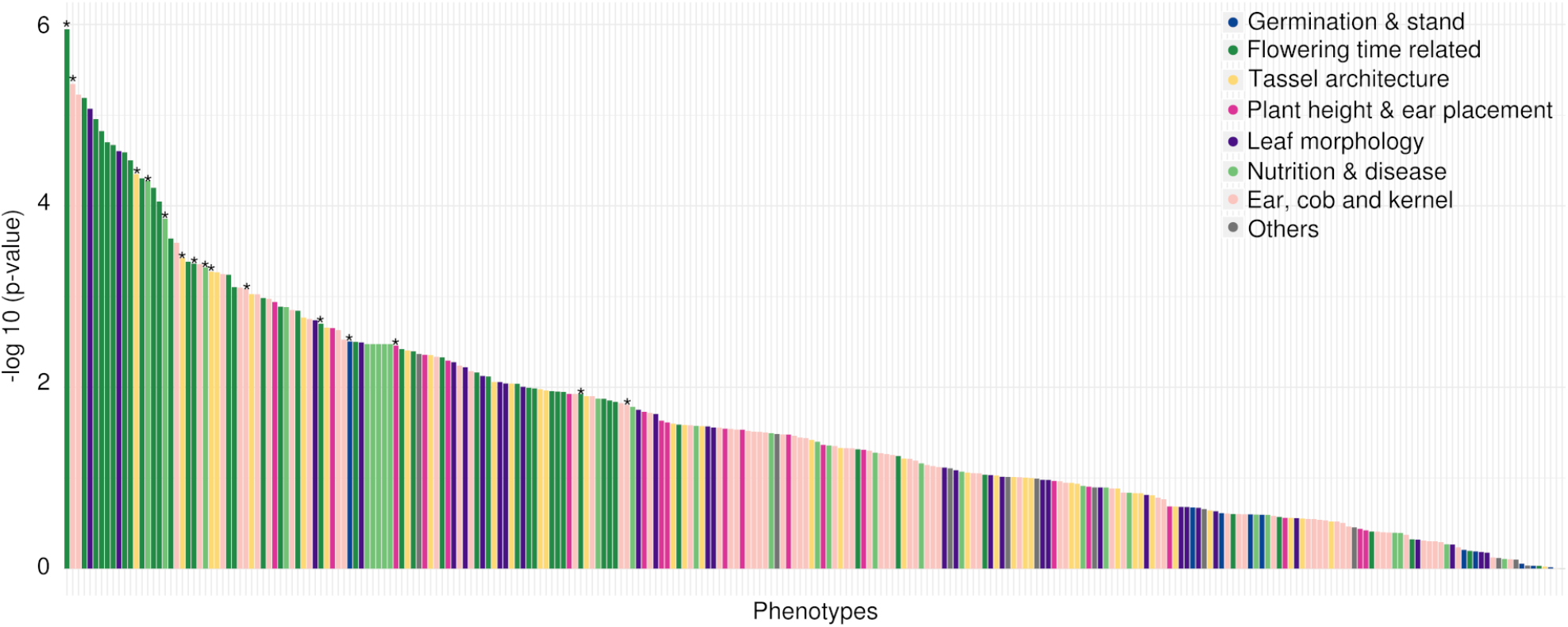
P-values assigned to the link between *an1* and individual phenotypes when the multi-SNP GPWAS algorithm was run with a single iteration and provided with only a single trait per analysis. Essentially this eliminates the effects of the forward selection and the “multi-trait” components of the GPWAS algorithm, retaining only the “multi-marker” components. Traits are sorted from most significant single-trait GPWAS association to least. Stars indicate those phenotypes which were associated with *an1* when all 260 phenotypes were analyzed together using the full GPWAS model.

**Figure S8.**
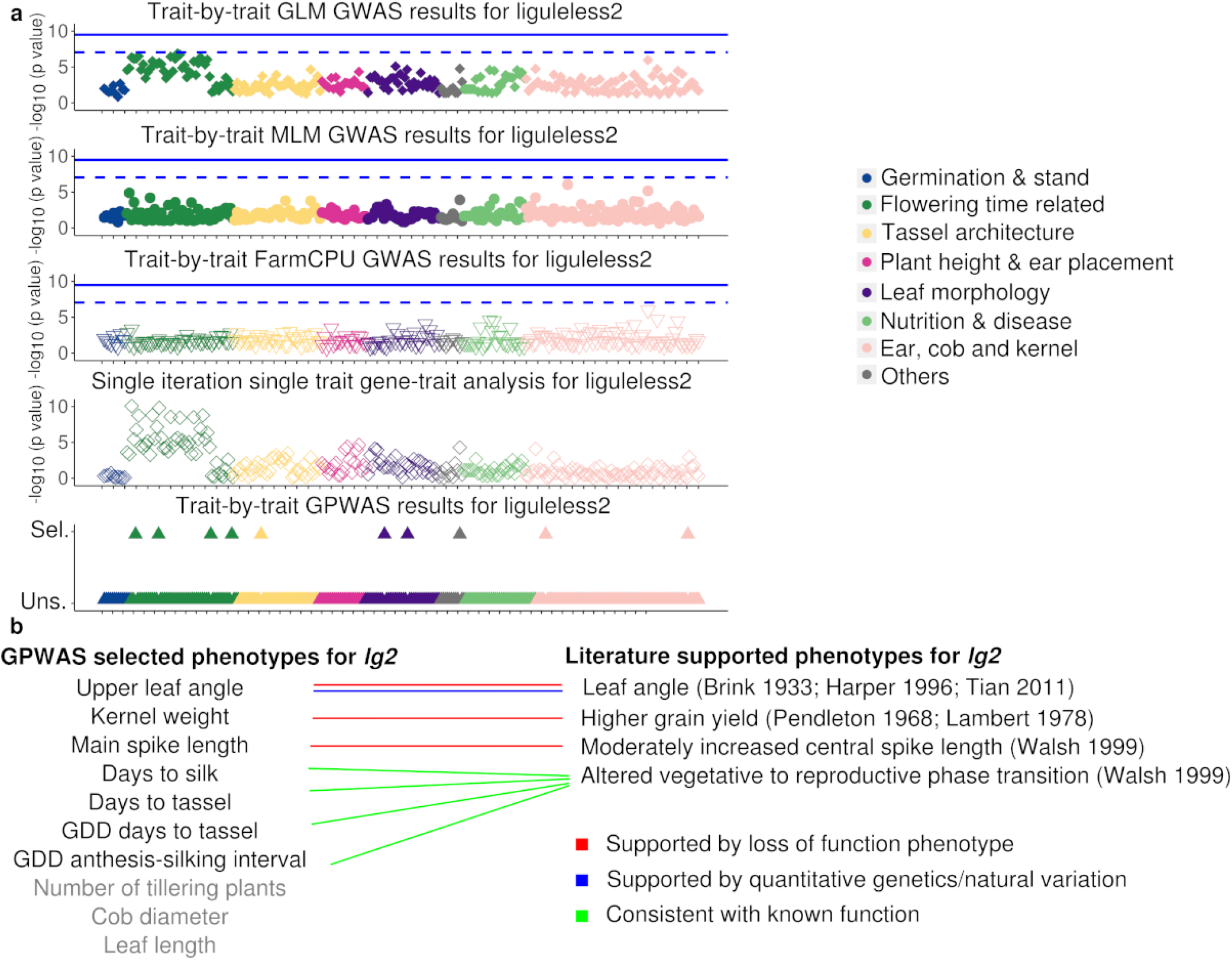
Evaluation of GLM GWAS, MLM GWAS, FarmCPU GWAS, single iteration single trait gene-trait analysis and GPWAS using the known maize gene *liguleless2* (*lg2*) (Zm00001d042777). (a) The dashed lines indicates a p value corresponding to 0.05 after a Bonferroni correction for independent tests on 557,968 (SNPs). Solid lines indicate the stricter multiple testing corrected threshold which considers both the number of SNPs and the number of phenotypes tested. In the GPWAS panel, Sel. and Uns. indicate traits that were selected and unselected respectively, in the model GPWAS fit for this particular gene. Phenotypes are ordered along the x-axis in the same order used for Figure 1, with each tick mark indicating a distance of five phenotypes. Phenotypes incorporated in the GPWAS model for *lg2* were as follows: days to silk (Summer 2006, Johnston, NC), days to tassel (Winter 2006, Ponce, PR), GDD days to tassel (Summer 2007, Champaign, IL), GDD anthesis-silking interval (Winter 2007, Miami-Dade, FL), main spike length (Summer 2006, Johnston, NC), leaf length (Summer 2006, Boone, MO), upper leaf angle (Summer 2006, Cayuga, NY), number of tillering plants (Summer 2007, Cayuga, NY), cob diameter (Winter 2006, Ponce, PR) and kernel weight (Summer 2007, Cayuga, NY). (b) The potential correspondence between phenotypes selected by the GPWAS model for *lg2*, and phenotypes either reported for loss of function *lg2* mutants or previous quantitative genetic analyses^36, 47, 49–52^.

**Figure S9.**
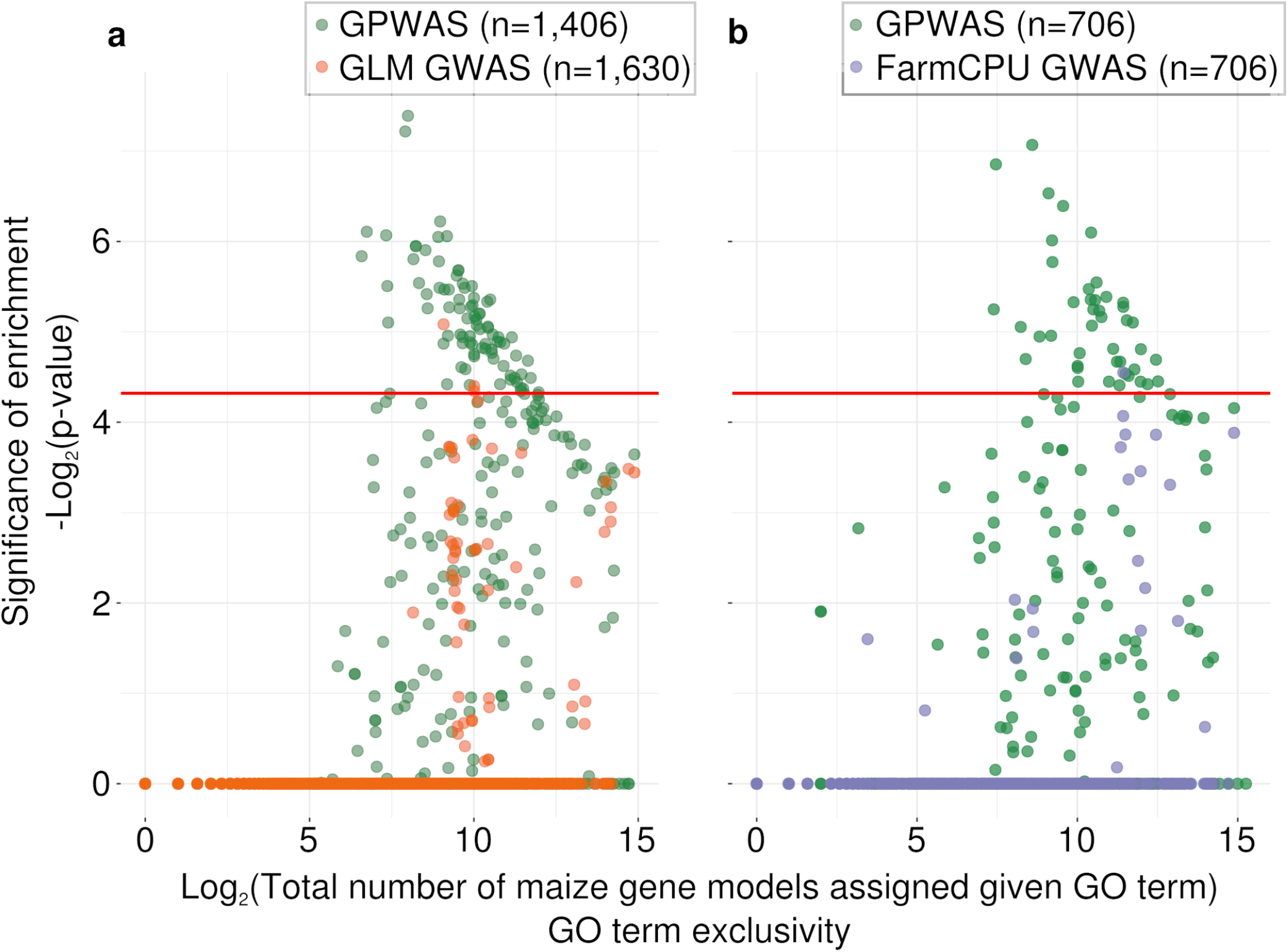
Comparison of GO enrichment/purification among genes uniquely identified as being associated with phenotypic variation using different statistical approaches. Each circle represents a single GO term in a single analysis. The position of each circle on the x axis indicates the total number of maize gene models which were assigned to this GO term in the maize GAMER dataset^96^. The position of each circle on the y-axis indicates the statistical significance of the enrichment or purification of this GO term in the given gene population relative to the background set of all annotated maize gene models. Red lines indicate the threshold for determining a significant GO term after a Bonferroni correction. (a) Comparison of the patterns of GO term enrichment/purification among genes either uniquely identified as being associated with phenotypic variation using a GLM GWAS analysis or uniquely identified as being associated with phenotypic variation in a GPWAS analysis. (b) As in panel a, but the comparison is between genes uniquely identified as being associated with phenotypic variation using a FarmCPU analysis or uniquely identified as being associated with phenotypic variation in a GPWAS analysis. Only the 706 genes uniquely identified using GPWAS with the strongest statistical signal were employed in panel b, to prevent any bias towards more significant p-values resulting from an analysis using a larger population of genes identified using GPWAS than those identified using FarmCPU.

**Figure S10.**
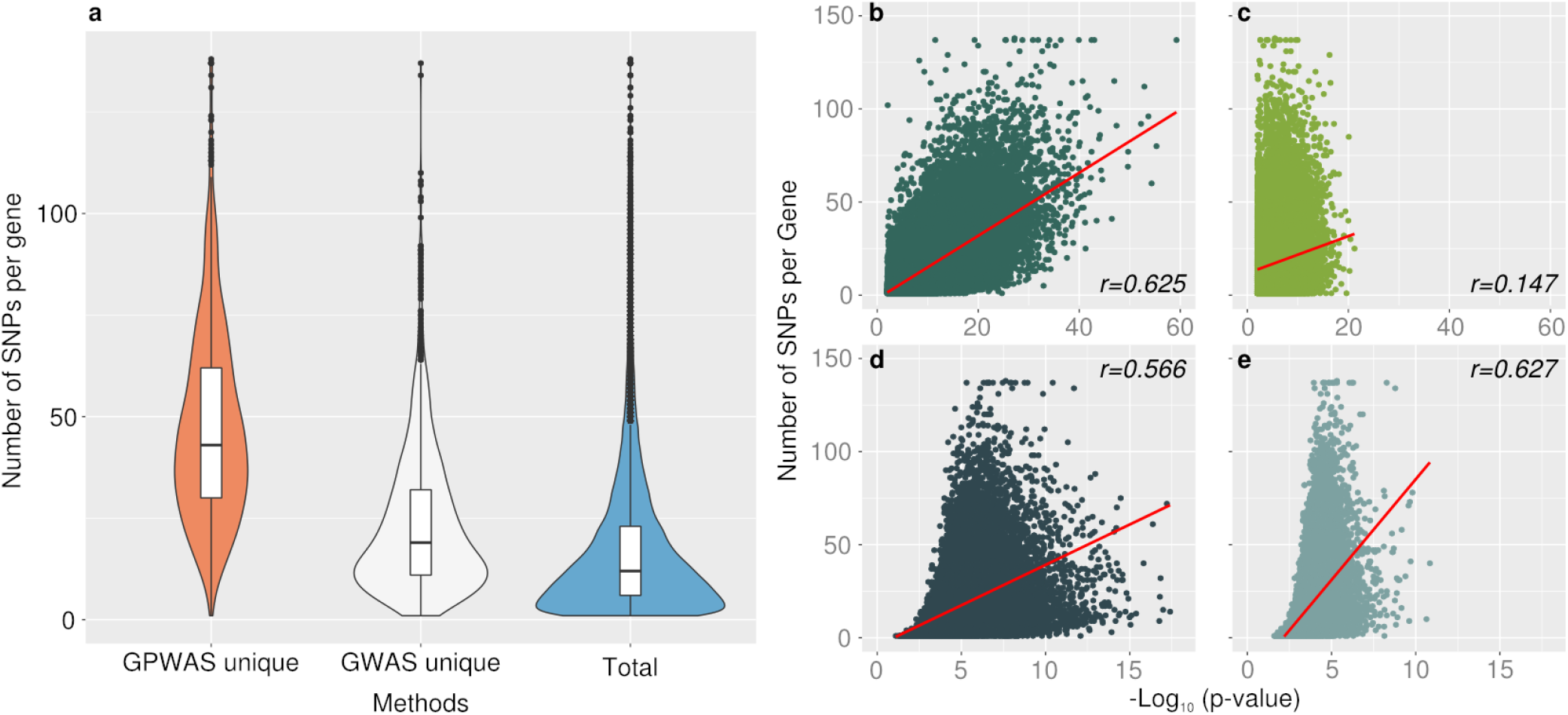
Number of SNPs identified per gene and the p-value of genes identified using different models. (a) The number of SNPs assigned to genes uniquely identified using either GPWAS or GLM GWAS, as well as the total number of genes with identified SNPs. SNPs assigned to gene regions were filtered and employed in all analyses. The maximum remaining number of SNPs per gene was 138. The distributions of the genes uniquely identified using GLM GWAS or GPWAS were statistically significantly different, p *<* 2.2e–16 (Mann-Whitney U test; two-sided). (b) Correlations between the SNP number per gene and the –log10 p-value of the total number of genes identified using GPWAS on real phenotype data. (c) Correlations between the SNP number per gene and the –log10 p-value of the total genes identified using GPWAS on randomly selected phenotype data from 20 permutations. (d) Correlations between the SNP number per gene and the –log10 p-value of the total genes identified using GLM GWAS on real phenotype data. (e) Correlations between the SNP number per gene and –log10 p-value of total genes identified using GLM GWAS on randomly selected phenotype data from 20 permutations. Spearman correlation methods were employed for the correlation test between SNP number and –log10 transformed p-value for each gene. Full statistical reports are presented in Supplementary Table 8.

**Figure S11.**
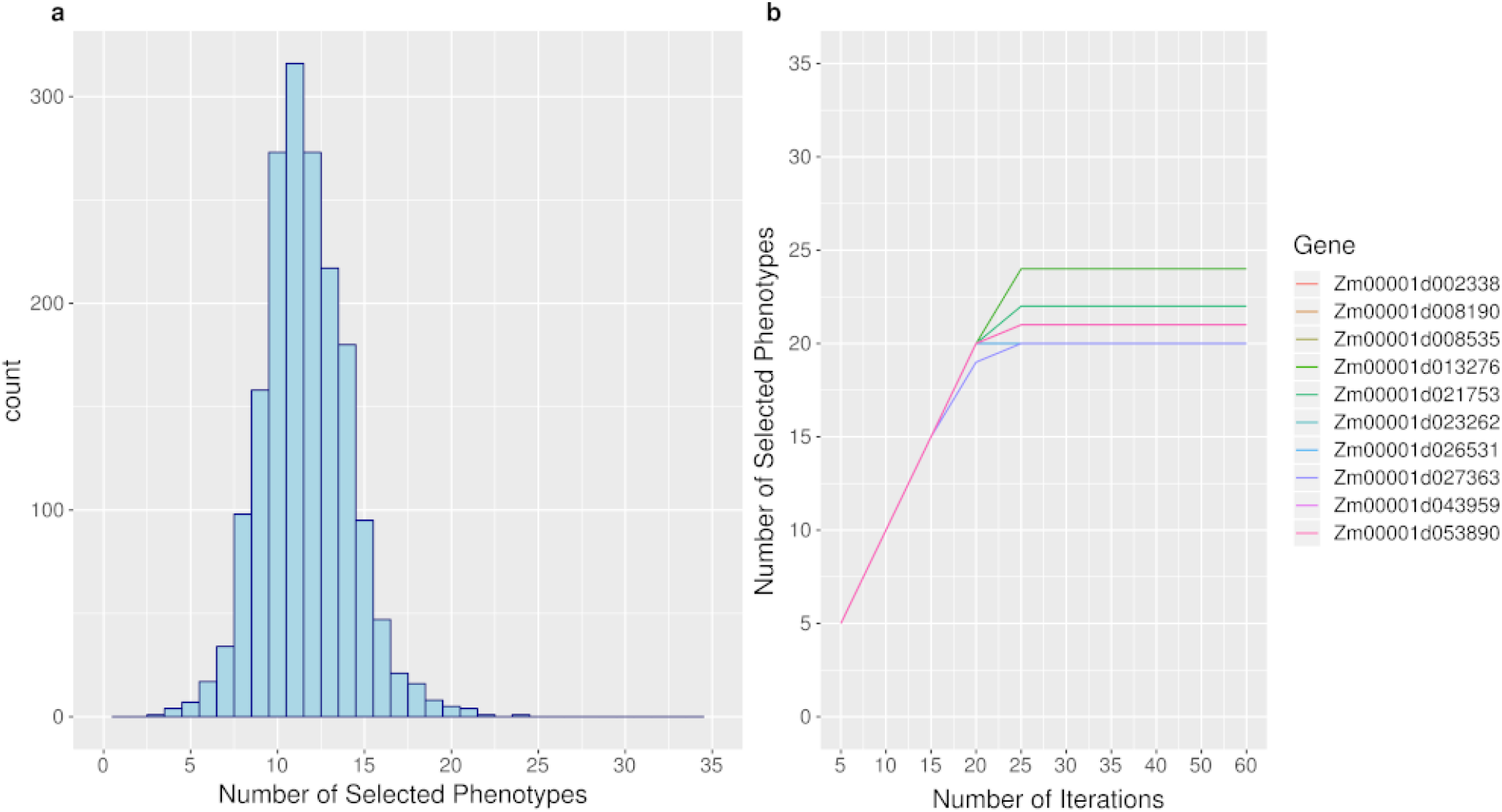
Estimating the number of iterations required for convergence of the forward selection model for GPWAS using the set of phenotypes and genotypic data for the maize 282 buckler goodman association panel employed here. (a) Distribution of the number of phenotypes incorporated into GPWAS models for the 1,776 genes identified as significantly linked to maize phenomic variation in this study; (b) Change in the number of phenotypes incorporated into genetic models as the number of iterations employed for GPWAS increases. Data shown for the 10 genes with the largest total number of phenotypes incorporated into their models among the 1,776 genes identified in this study.

## References

1. Schnable, J. C. & Freeling, M. Genes identified by visible mutant phenotypes show increased bias toward one of two subgenomes of maize. PloS one 6, e17855 (2011).

2. Lamesch, P. et al. The arabidopsis information resource (tair): improved gene annotation and new tools. Nucleic acids research 40, D1202–D1210 (2011).

3. Rhee, S. Y. & Mutwil, M. Towards revealing the functions of all genes in plants. Trends plant science 19, 212–221 (2014).

4. Schofield, P. N., Hoehndorf, R. & Gkoutos, G. V. Mouse genetic and phenotypic resources for human genetics. Hum. mutation 33, 826–836 (2012).

5. Chong, J. X. et al. The genetic basis of mendelian phenotypes: discoveries, challenges, and opportunities. The Am. J. Hum. Genet. 97, 199–215 (2015).

6. Schnable, J. C. Genes and gene models, an important distinction. New Phytol. (2019).

7. Lock, A. et al. Pombase 2018: user-driven reimplementation of the fission yeast database provides rapid and intuitive access to diverse, interconnected information. Nucleic acids research 47, D821–D827 (2018).

8. Sax, K. The association of size differences with seed-coat pattern and pigmentation in phaseolus vulgaris. Genetics 8, 552 (1923).

9. Sprague, G. The location of dominant favorable genes in maize by means of an inversion. Genetics 26, 143–149 (1941).

10. Atwell, S. et al. Genome-wide association study of 107 phenotypes in arabidopsis thaliana inbred lines. Nature 465, 627 (2010).

11. Klein, R. J. et al. Complement factor h polymorphism in age-related macular degeneration. Science 308, 385–389 (2005).

12. DeWan, A. et al. Htra1 promoter polymorphism in wet age-related macular degeneration. Science 314, 989–992 (2006).

13. Korte, A. et al. A mixed-model approach for genome-wide association studies of correlated traits in structured populations. Nat. genetics 44, 1066 (2012).

14. O’Reilly, P. F. et al. Multiphen: joint model of multiple phenotypes can increase discovery in gwas. PloS one 7, e34861 (2012).

15. Van der Sluis, S., Posthuma, D. & Dolan, C. V. Tates: efficient multivariate genotype-phenotype analysis for genome-wide association studies. PLoS genetics 9, e1003235 (2013).

16. Stephens, M. A unified framework for association analysis with multiple related phenotypes. PloS one 8, e65245 (2013).

17. Zhou, X. & Stephens, M. Genome-wide efficient mixed-model analysis for association studies. Nat. genetics 44, 821 (2012).

18. Wang, Y. et al. Pleiotropy analysis of quantitative traits at gene level by multivariate functional linear models. Genet. epidemiology 39, 259–275 (2015).

19. Turley, P. et al. Multi-trait analysis of genome-wide association summary statistics using mtag. Nat. genetics 50, 229 (2018).

20. Pitchers, W. et al. A multivariate genome-wide association study of wing shape in drosophila melanogaster. Genetics genetics–301342 (2019).

21. Lin, H.-y. et al. Substantial contribution of genetic variation in the expression of transcription factors to phenotypic variation revealed by erd-gwas. Genome biology 18, 192 (2017).

22. Kremling, K., Diepenbrock, C., Gore, M., Buckler, E. & Bandillo, N. Transcriptome-wide association supplements genome-wide association in zea mays. bioRxiv 363242 (2018).

23. Wen, W. et al. Metabolome-based genome-wide association study of maize kernel leads to novel biochemical insights. Nat. communications 5, 3438 (2014).

24. Matsuda, F. et al. Metabolome-genome-wide association study dissects genetic architecture for generating natural variation in rice secondary metabolism. The Plant J. 81, 13–23 (2015).

25. Diepenbrock, C. H. et al. Novel loci underlie natural variation in vitamin e levels in maize grain. The Plant Cell tpc–00475 (2017).

26. Kremling, K. A. et al. Dysregulation of expression correlates with rare-allele burden and fitness loss in maize. Nature 555, 520 (2018).

27. Walter, A., Liebisch, F. & Hund, A. Plant phenotyping: from bean weighing to image analysis. Plant methods 11, 14 (2015).

28. Araus, J. L., Kefauver, S. C., Zaman-Allah, M., Olsen, M. S. & Cairns, J. E. Translating high-throughput phenotyping into genetic gain. Trends plant science 23, 451–466 (2018).

29. Flint-Garcia, S. A. et al. Maize association population: a high-resolution platform for quantitative trait locus dissection. The Plant J. 44, 1054–1064 (2005).

30. Bukowski, R. et al. Construction of the third-generation zea mays haplotype map. GigaScience 7, gix134 (2017).

31. Wallace, J. G. et al. Association mapping across numerous traits reveals patterns of functional variation in maize. PLoS genetics 10, e1004845 (2014).

32. Zhao, W. et al. Panzea: a database and resource for molecular and functional diversity in the maize genome. Nucleic Acids Res. 34, D752–D757 (2006).

33. Dahl, A. et al. A multiple-phenotype imputation method for genetic studies. Nat. genetics 47, 466 (2015).

34. Consortium, I. H. et al. A haplotype map of the human genome. Nature 437, 1299 (2005).

35. Li, J. & Ji, L. Adjusting multiple testing in multilocus analyses using the eigenvalues of a correlation matrix. Heredity 95, 221 (2005).

36. Tian, F. et al. Genome-wide association study of leaf architecture in the maize nested association mapping population. Nat. genetics 43, 159 (2011).

37. Namjou, B. et al. Phenome-wide association study (phewas) in emr-linked pediatric cohorts. Front. genetics 5, 401 (2014).

38. Price, A. L. et al. Principal components analysis corrects for stratification in genome-wide association studies. Nat. genetics 38, 904 (2006).

39. Yu, J. et al. A unified mixed-model method for association mapping that accounts for multiple levels of relatedness. Nat. genetics 38, 203 (2006).

40. Liu, X., Huang, M., Fan, B., Buckler, E. S. & Zhang, Z. Iterative usage of fixed and random effect models for powerful and efficient genome-wide association studies. PLoS genetics 12, e1005767 (2016).

41. McMullen, M. D. et al. Genetic properties of the maize nested association mapping population. Science 325, 737–740 (2009).

42. Owens, B. F. et al. A foundation for provitamin a biofortification of maize: genome-wide association and genomic prediction models of carotenoid levels. Genetics 198, 1699–1716 (2014).

43. Bensen, R. J. et al. Cloning and characterization of the maize an1 gene. The Plant Cell 7, 75–84 (1995).

44. Brown, P. J. et al. Distinct genetic architectures for male and female inflorescence traits of maize. PLoS genetics 7, e1002383 (2011).

45. Peng, J. & Harberd, N. P. The role of ga-mediated signalling in the control of seed germination. Curr. opinion plant biology 5, 376–381 (2002).

46. Landoni, M. et al. The an1-4736 mutation of anther ear1 in maize alters scotomorphogenesis and the light response. Plant science 172, 172–180 (2007).

47. Brink, R. Heritable characters in maize: Xlvi—ligu1eless-2. J. Hered. 24, 325–326 (1933).

48. Walsh, J., Waters, C. A. & Freeling, M. The maize geneliguleless2 encodes a basic leucine zipper protein involved in the establishment of the leaf blade–sheath boundary. Genes & Dev. 12, 208–218 (1998).

49. Harper, L. & Freeling, M. Interactions of liguleless1 and liguleless2 function during ligule induction in maize. Genetics 144, 1871–1882 (1996).

50. Pendleton, J., Smith, G., Winter, S. & Johnston, T. Field investigations of the relationships of leaf angle in corn (zea mays l.) to grain yield and apparent photosynthesis 1. Agron. J. 60, 422–424 (1968).

51. Lambert, R. & Johnson, R. Leaf angle, tassel morphology, and the performance of maize hybrids 1. Crop. Sci. 18, 499–502 (1978).

52. Walsh, J. & Freeling, M. The liguleless2 gene of maize functions during the transition from the vegetative to the reproductive shoot apex. The Plant J. 19, 489–495 (1999).

53. Li, M., Zhong, W., Yang, F. & Zhang, Z. Genetic and molecular mechanisms of quantitative trait loci controlling maize inflorescence architecture. Plant Cell Physiol. 59, 448–457 (2018).

54. Liu, L. et al. Krn4 controls quantitative variation in maize kernel row number. PLoS genetics 11, e1005670 (2015).

55. Paz-Ares, J., Ghosal, D., Wienand, U., Peterson, P. & Saedler, H. The regulatory c1 locus of zea mays encodes a protein with homology to myb proto-oncogene products and with structural similarities to transcriptional activators. The EMBO J. 6, 3553–3558 (1987).

56. Xu, X. et al. Sequence analysis of the cloned glossy8 gene of maize suggests that it may code for a [beta]-ketoacyl reductase required for the biosynthesis of cuticular waxes. Plant physiology 115, 501–510 (1997).

57. Chuck, G., Meeley, R. B. & Hake, S. The control of maize spikelet meristem fate by theapetala2-like gene indeterminate spikelet1. Genes & Dev. 12, 1145–1154 (1998).

58. Sturaro, M. et al. Cloning and characterization of glossy1, a maize gene involved in cuticle membrane and wax production. Plant physiology 138, 478–489 (2005).

59. Stelpflug, S. C. et al. An expanded maize gene expression atlas based on rna sequencing and its use to explore root development. The plant genome 9 (2016).

60. Morris, G. P. et al. Population genomic and genome-wide association studies of agroclimatic traits in sorghum. Proc. Natl. Acad. Sci. 110, 453–458 (2013).

61. Huang, X. et al. Genome-wide association studies of 14 agronomic traits in rice landraces. Nat. genetics 42, 961 (2010).

62. Rencher, A. C. & Schaalje, G. B. Linear models in statistics (John Wiley & Sons, 2008).

63. Piepho, H., Möhring, J., Melchinger, A. & Büchse, A. Blup for phenotypic selection in plant breeding and variety testing. Euphytica 161, 209–228 (2008).

64. Kusmec, A., Srinivasan, S., Nettleton, D. & Schnable, P. S. Distinct genetic architectures for phenotype means and plasticities in zea mays. Nat. plants 3, 715 (2017).

65. Rodgers-Melnick, E., Vera, D. L., Bass, H. W. & Buckler, E. S. Open chromatin reveals the functional maize genome. Proc. Natl. Acad. Sci. 113, E3177–E3184 (2016).

66. Studer, A., Zhao, Q., Ross-Ibarra, J. & Doebley, J. Identification of a functional transposon insertion in the maize domestication gene tb1. Nat. genetics 43, 1160 (2011).

67. Castelletti, S., Tuberosa, R., Pindo, M. & Salvi, S. A mite transposon insertion is associated with differential methylation at the maize flowering time qtl vgt1. G3: Genes, Genomes, Genet. 4, 805–812 (2014).

68. Remington, D. L. et al. Structure of linkage disequilibrium and phenotypic associations in the maize genome. Proc. Natl. Acad. Sci. 98, 11479–11484 (2001).

69. Romay, M. C. et al. Comprehensive genotyping of the usa national maize inbred seed bank. Genome biology 14, R55 (2013).

70. Zhang, W. et al. High-resolution mapping of open chromatin in the rice genome. Genome research 22, 151–162 (2012).

71. Turco, G., Schnable, J. C., Pedersen, B. & Freeling, M. Automated conserved non-coding sequence (cns) discovery reveals differences in gene content and promoter evolution among grasses. Front. plant science 4, 170 (2013).

72. Oka, R. et al. Genome-wide mapping of transcriptional enhancer candidates using dna and chromatin features in maize. Genome biology 18, 137 (2017).

73. Lai, X. et al. Stag-cns: An order-aware conserved noncoding sequences discovery tool for arbitrary numbers of species. Mol. plant 10, 990–999 (2017).

74. Lu, Z., Ricci, W. A., Schmitz, R. J. & Zhang, X. Identification of cis-regulatory elements by chromatin structure. Curr. opinion plant biology 42, 90–94 (2018).

75. Lloyd, J. P., Tsai, Z. T.-Y., Sowers, R. P., Panchy, N. L. & Shiu, S.-H. A model-based approach for identifying functional intergenic transcribed regions and noncoding rnas. Mol. biology evolution 35, 1422–1436 (2018).

76. Bennetzen, J. L., Coleman, C., Liu, R., Ma, J. & Ramakrishna, W. Consistent over-estimation of gene number in complex plant genomes. Curr. opinion plant biology 7, 732–736 (2004).

77. Gerstein, M. B. et al. What is a gene, post-encode? history and updated definition. Genome research 17, 669–681 (2007).

78. Schnable, J. C. Genome evolution in maize: from genomes back to genes. Annu. Rev. Plant Biol. 66, 329–343 (2015).

79. Browning, B. L. & Browning, S. R. Genotype imputation with millions of reference samples. The Am. J. Hum. Genet. 98, 116–126 (2016).

80. Listgarten, J. et al. Improved linear mixed models for genome-wide association studies. Nat. methods 9, 525 (2012).

81. Rincent, R. et al. Recovering power in association mapping panels with variable levels of linkage disequilibrium. Genetics 197, 375–387 (2014).

82. Draper, N. R. & Smith, H. Applied regression analysis, vol. 326 (John Wiley & Sons, 1998).

83. Johnson, R. A., Wichern, D. W. et al. Applied multivariate statistical analysis, vol. 5 (Prentice hall Upper Saddle River, NJ, 2002).

84. Lippert, C. et al. Fast linear mixed models for genome-wide association studies. Nat. methods 8, 833 (2011).

85. Zhou, X. A unified framework for variance component estimation with summary statistics in genome-wide association studies. The annals applied statistics 11, 2027 (2017).

86. Schaeffer, M. L. et al. Maizegdb: curation and outreach go hand-in-hand. Database 2011 (2011).

87. Bolger, A. M., Lohse, M. & Usadel, B. Trimmomatic: a flexible trimmer for illumina sequence data. Bioinformatics 30, 2114–2120 (2014).

88. Wu, T. D. & Nacu, S. Fast and snp-tolerant detection of complex variants and splicing in short reads. Bioinformatics 26, 873–881 (2010).

89. Li, H. et al. The sequence alignment/map format and samtools. Bioinformatics 25, 2078–2079 (2009).

90. Trapnell, C. et al. Differential gene and transcript expression analysis of rna-seq experiments with tophat and cufflinks. Nat. protocols 7, 562 (2012).

91. Schnable, J. C. Sorghum version 3, maize versions 3 and 4 syntenic gene list. FigShare.

92. McCormick, R. F. et al. The sorghum bicolor reference genome: improved assembly, gene annotations, a transcriptome atlas, and signatures of genome organization. The Plant J. 93, 338–354 (2018).

93. Bennetzen, J. L. et al. Reference genome sequence of the model plant setaria. Nat. biotechnology 30, 555 (2012).

94. Zhang, Y. et al. Differentially regulated orthologs in sorghum and the subgenomes of maize. The Plant Cell tpc–00354 (2017).

95. Brohammer, A. B., Kono, T. J., Springer, N. M., McGaugh, S. E. & Hirsch, C. N. The limited role of differential fractionation in genome content variation and function in maize (zea mays l.) inbred lines. The Plant J. 93, 131–141 (2018).

96. Wimalanathan, K., Friedberg, I., Andorf, C. M. & Lawrence-Dill, C. J. Maize go annotation—methods, evaluation, and review (maize-gamer). Plant Direct 2, e00052 (2018).

97. Klopfenstein, D. et al. Goatools: A python library for gene ontology analyses. Sci. reports 8, 10872 (2018).

98. Yang, J., Lee, S. H., Goddard, M. E. & Visscher, P. M. Gcta: a tool for genome-wide complex trait analysis. The Am. J. Hum. Genet. 88, 76–82 (2011).

